# Providing insight into the mechanism of action of Cationic Lipidated Oligomers (CLOs) using metabolomics

**DOI:** 10.1101/2024.02.19.581110

**Authors:** Maytham Hussein, Muhammad Bilal Hassan Mahboob, Jessica R. Tait, James L. Grace, Véronique Montembault, Laurent Fontaine, John F. Quinn, Tony Velkov, Michael R. Whittaker, Cornelia B. Landersdorfer

## Abstract

The increasing resistance of clinically relevant microbes against current commercially available antimicrobials underpins the urgent need for alternative and novel treatment strategies. Cationic lipidated oligomers (CLOs) are innovative alternatives to antimicrobial peptides, and have reported antimicrobial potential. An understanding of their antimicrobial mechanism of action is required to rationally design future treatment strategies for CLOs, either in monotherapy or synergistic combinations. In the present study, metabolomics was used to investigate the potential metabolic pathways involved in the mechanisms of antibacterial activity of one CLO, C_12_-o-(BG-D)-10, which we have previously shown to be effective against methicillin-resistant *Staphylococcus aureus* (MRSA) ATCC 43300. The metabolomes of MRSA ATCC 43300 at 1, 3 and 6 h following treatment with C_12_-o-(BG-D)-10 (48 µg/mL i.e., 3x MIC) were compared to those of the untreated controls.

Our findings reveal that the studied CLO, C_12_-o-(BG-D)-10, disorganized the bacterial membrane as the first step towards its antimicrobial effect, as evidenced by marked perturbations in the bacterial membrane lipids and peptidoglycan biosynthesis observed at early time points i.e., 1, and 3 h. Central carbon metabolism, and biosynthesis of DNA, RNA, and arginine were also vigorously perturbed, mainly at early time points. Moreover, bacterial cells were under osmotic and oxidative stress across all time points, evident by perturbations of trehalose biosynthesis and pentose phosphate shunt. Overall, this metabolomics study has, for the first time, revealed that the antimicrobial action of C_12_-o-(BG-D)-10 may potentially stem from the dysregulation of multiple metabolic pathways.

**Importance:** Antimicrobial resistance poses a significant challenge to healthcare systems worldwide. Novel anti-infective therapeutics are urgently needed to combat drug-resistant microorganisms. Cationic lipidated oligomers (CLOs) show promise as new antibacterial agents against Gram-positive pathogens like *Staphylococcus aureus* (MRSA). Understanding their molecular mechanism(s) of antimicrobial action may help design synergistic CLO treatments along with monotherapy. Here, we describe the first metabolomics study to investigate the killing mechanism(s) of CLOs against MRSA. The results of our study indicate that the CLO, C_12_-o-(BG-D)-10, had a notable impact on the biosynthesis and organization of the bacterial cell envelope. C_12_-o-(BG-D)-10 also inhibits arginine, histidine, central carbon metabolism, and trehalose production, adding to its antibacterial characteristics. This work illuminates the unique mechanism of action of C_12_-o-(BG-D)-10 and opens an avenue to design innovative antibacterial oligomers/polymers for future clinical applications.

## INTRODUCTION

Antimicrobial resistance (AMR) represents a pressing global health challenge, posing significant threats to successful treatment of infections and patient outcomes [1]. Common mechanisms through which microbes develop resistance to antimicrobials include reduced drug uptake, drug target modification, modification of drug and drug efflux; these may emerge due to genetic mutations or horizontal gene transfer [2]. Emergence of AMR is largely attributed to the inappropriate and excessive use of various antibacterial agents, both within the healthcare sector and the agricultural industry [3]. The dwindling arsenal of effective treatment options poses a substantial challenge, jeopardizing our ability to combat infections successfully. As a direct consequence, mortality rates are rising, as once-treatable infections become increasingly difficult to treat. Beyond the immediate health impacts, AMR causes substantial economic burden through heightened healthcare costs, including due to prolonged hospital stays, higher likelihood of readmission and increased expenditure on additional treatments [4]. Recognizing AMR as a critical issue, global health organizations such as the Infectious Diseases Society of America in 2009, the World Health Organization (WHO) in 2017, and the Centers for Disease Control and Prevention (US-CDC) of the United States in 2019, have emphasized the need for multifaceted strategies, including prudent antimicrobial use, surveillance, and the development of novel drugs. They also issued lists of critical pathogens on which to focus the research and development of new antimicrobials [5–8].

While conventional antimicrobials are currently used for treating infections, the rapid increase in AMR suggests the need for alternative treatments [9]. Antimicrobial peptides (AMPs) have been investigated and have been demonstrated to be very effective in the killing of microbes [10–12]. However, AMPs have limitations, such as instability, toxicity, and high production costs [13, 14]. Recently, synthetic analogs of AMPs, i.e. antimicrobial polymers that mimic the structural features (cationic and hydrophobic moieties) of naturally occurring AMPs, have been designed and shown to overcome AMP limitations, in *in vitro* studies and *in vivo* mouse models [15]. This class of molecules has several advantages, such as broad-spectrum antimicrobial activities, a low tendency for resistance development and a rapid bactericidal effect [11, 16–22]. Furthermore, antimicrobial polymers offer extra advantages compared to AMPs in terms of stability, durability, and ease of large-scale production [17]. These antimicrobial polymers have a broad range of mechanisms of actions from membrane disruption to intracellular target inhibition, depending on the structural features [17]. However, the full mechanism(s) of action of antimicrobial polymers remains unclear in the literature as detailed structure-property relationships are difficult to elucidate.

*Staphylococcus aureus* is a Gram-positive bacterial species categorized as a high-priority pathogen by the WHO and a serious threat by the US-CDC [23, 24]. *S. aureus* is part of the normal flora of humans and usually does not cause infection while on the skin. However, when the skin barrier is damaged, *S. aureus* may enter the underlying tissue or bloodstream and cause a wide variety of infections [25]. Currently, methicillin-resistant *S. aureus* (MRSA) is one of the main nosocomial pathogens and is prevalent in hospital settings [26]. In 2019, 473,000 deaths were associated with MRSA infections globally [27]. In Australia, MRSA infections were associated with increased inpatient mortality, as well as greater expense and longer hospital length of stay compared with methicillin-susceptible *S. aureus* [28].

In recent work, our group has shown that cationic lipidated oligomers (CLOs) demonstrate structure-dependent antimicrobial activity against both Gram-positive and Gram-negative bacteria and fungi [29]. Interestingly, while these CLOs have been designed with the same structural features (with cationic residues and lipid tail) as the clinically used AMP colistin [30], they appear to have wider applicability. The synthesized CLOs have repeating cationic residues [e.g. tertiary amine, primary amine (mimicking lysine), guanidine (mimicking arginine), or imidazole (mimicking histidine)], which help these CLOs to electrostatically bind to the negatively charged bacterial membranes [17]. The lipid tail in each of the CLOs helps to disrupt bacterial membranes more effectively [16]. One CLO in particular, i.e. C_12_-o-(BG-D)-10, with ten cationic guanidine residues and a C_12_ hydrocarbon lipid tail, exhibited marked antimicrobial activity against MRSA [29]. These guanidine groups have been reported to form multidentate hydrogen bonds with sulfate and phosphate heads on the bacterial anionic membranes, leading to efficient bacterial membrane integration [31–33]. However, their precise mechanism/s of antimicrobial action remain largely unknown to date.

Metabolomics has emerged as a critical tool for the elucidation of the mechanisms of action of AMPs such as polymyxins [34, 35]. In this study, we shine a light on the mechanism(s) of bacterial killing by the antimicrobial CLO, C_12_-o-(BG-D)-10, against MRSA strain ATCC 43300 using untargeted metabolomics.

## RESULTS AND DISCUSSION

The antibacterial activity of C_12_-o-(BG-D)-10 was previously assessed against *S. aureus* ATCC 43300 and demonstrated an appreciable activity with a MIC of 8-16 µg/mL [29]. Additionally, C_12_-o-(BG-D)-10 exhibited an interesting mechanism of action when comparing dose-dependent membrane disruption (*via* fluorescence Propidium iodide (PI) assay) and growth inhibition **(Fig. S1)**. Specifically, C_12_-o-(BG-D)-10 exhibited only minor interaction with the bacterial membrane, with only ∼30% membrane damage observed (relative to a melittin control) at the highest concentration tested. These results suggested that there could be potentially a secondary mechanism contributing to the observed antimicrobial activity for C_12_-o-(BG-D)-10 [29]. To interrogate this complex mode of action, a metabolomics study was performed using an initial inoculum of 10^8^ colony forming units (CFU)/mL with samples at 1, 3, and 6 h. The 48 µg/mL (3x MIC) C_12_-o-(BG-D)-10 concentration provided maximal bacterial killing at 1 h with ∼1.5 log_10_CFU/mL decrease compared to the control **(Fig. S2a)**. Metabolomics results of different perturbed metabolic pathways of MRSA are discussed below.

### Multivariate and univariate analysis

A total of 1578 putative metabolites were identified under all treatment conditions. Out of these, 33% were not mapped to known metabolic pathways and 67% were mapped to known metabolic pathways according to common databases, e.g. PseudoCyc, MetaCyc, and LipidMaps databases. Most of the acquired metabolites belonged to lipid (18%), peptide (18%), and amino-acid (17%) metabolism, while the minority of metabolites belonged to carbohydrate (6%), nucleotide (4%), secondary metabolites (2%), energy (1%), and glycan (0.5%) metabolite classes. The same databases were then used to designate metabolic classes or map the unmapped metabolites. A univariate data analysis was performed using two-sample t-tests [≥1-log_2_-fold change (FC); *t*-tests, False Discovery Rate (FDR) adjusted *p*-value ≤0.05] to determine significantly perturbed metabolites across all time points (1, 3 and 6 h). This analysis identified ∼476 significantly perturbed metabolites [295, 241 and 185 at 1, 3 and 6 h, respectively] **(Fig. S3)**. Across all time points, there were 53 overlapping metabolites with 155, 72, and 57 unique metabolites at 1, 3, and 6 h, respectively **(Fig. S3)**. The majority of these metabolites were diminished in response to treatment with C_12_-o-(BG-D)-10. The heatmap showed that the intensities of metabolites varied after treatment with C_12_-o-(BG-D)-10 across all time points especially at 1 h **(Fig. S4).** The reproducibility for all sample groups was acceptable across all time points (1, 3 and 6 h), where the median relative standard deviations (RSDs) across all time points were 14% to 16% for untreated (control) groups, 17% to 22% for treated samples and 12% for the QC group, consistent with some baseline variability in the dynamics of ordinary bacterial metabolism with and without C_12_-o-(BG-D)-10 treatment **(Table 1)**. The well-separated treatment and control groups in the principal component analysis (PCA) revealed that C_12_-o-(BG-D)-10 treatment altered the metabolomic profile of MRSA across all time points **(Fig. S5)**. The classification of the significantly impacted metabolites across all time points revealed that the lipids, peptides, amino acids, and carbohydrate (including glycans) metabolites were largely impacted while the nucleotides metabolites and energy metabolite were less significantly perturbed across all the time points **(Fig. S6)**.

**Table 1:** Median relative standard deviations (RSDs) of all the metabolites of MRSA before and after treatment with C_12_-o-(BG-D)-10 across all the time points i.e., 1, 3, and 6 h.

| Median RSD% |  |  |  |  |
| --- | --- | --- | --- | --- |
|  | 1h | 3h | 6h | QC |
| **Control**<br>**(untreated)** | 14 | 16 | 16 | 12 |
| **Treatment**<br>**C<sub>12</sub>-o-(BG-D)-10** | 22 | 22 | 17 |  |

### Pathway enrichment analysis for the significantly perturbed metabolites

C_12_-o-(BG-D)-10 treatment induced extensive perturbations in the metabolomic profile of MRSA ATCC 43300 across all time points. Consequently, therefore we mapped and analyzed the significantly impacted metabolic features across all time points (i.e., 1, 3 and 6 h). The mapping of the significantly perturbed features of MRSA revealed that glycerophospholipids and FAs (fatty acids) metabolism, peptidoglycan and teichoic acid biosynthesis, DNA and RNA biosynthesis/nucleotide biosynthesis, central carbon metabolism, arginine biosynthesis, histidine metabolism, and pantothenate and Co-enzyme A (CoA) biosynthesis were among the most significantly perturbed pathways (Table S1).

### Glycerophospholipid and fatty acid metabolism

Following C_12_-o-(BG-D)-10 treatment, lipids were significantly perturbed, particularly the glycerophospholipids and fatty acids classes compared to other classes, i.e., glycerolipids, sphingolipids and sterol lipids. Among glycerophospholipids, long chain glycerophosphoglycerols, including PG(25:0), PG(19:0) and PG(14:0), were largely perturbed at 1 h **(Fig. 1a)**. The abundance of a highly important metabolite involved in the synthesis of bacterial membrane lipids and known precursor of teichoic acid in the Gram-positive bacterial cell wall, CDP-glycerol, was significantly decreased (log_2_FC = -3.01) at 1 h **(Fig. 1a)** [36]. The levels of two more well-known metabolites involved in bacterial membrane lipids biosynthesis, *sn*-glycerol 3-phosphate and *sn*-glycero-3-phosphocholine, were substantially decreased at 1 h post C_12_-o-(BG-D)-10 treatment (log_2_FC = -6.69 and -4.73, respectively) **(Fig. 1a)** [37–39]. Also, di-trans,poly-cis-undecaprenyl phosphate (also known lipid-P, bactoprenol and C55-P) was significantly perturbed (log_2_FC = 3.9) at 1 h; lipid-P, is involved in transporting peptidoglycan subunits from the cytoplasmic face of the cell membrane through the periplasmic space to the extracellular surface and plays a role in the synthesis of teichoic acid of the bacterial cell envelope [40–44]. Significant perturbations were also evident in the level of choline phosphate (log_2_FC = -3.0), which has a crucial role in the biosynthesis of wall teichoic acid in bacteria [45]. The abundance of (R)-3-hydroxybutanoate, an essential precursor involved in the synthesis of the bacterial membrane, was significantly decreased at 1 h (log_2_FC = -2.6) **(Fig. 1a)** [46].

**Figure 1.**
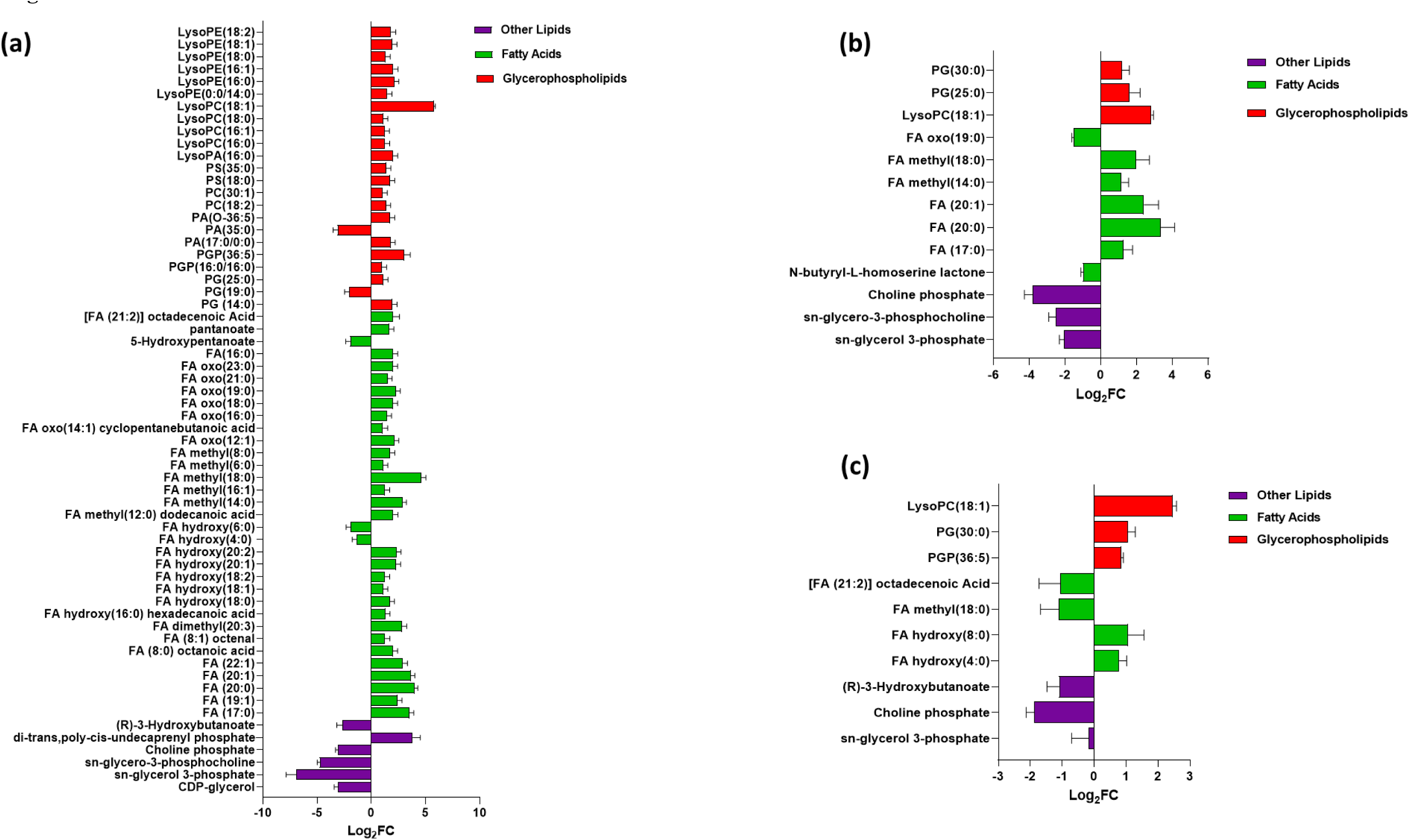
Significantly impacted lipids in MRSA ATCC 43300 following treatment with C_12_-o-(BG-D)-10 at 1 h (a), 3 h (b), and 6 h (c). Putative lipid names are assigned based on accurate mass (≥ 1.0-log_2_-FC; *p*<0.05). PE, phosphoethanolamines; PG, glycerophosphoglycerols; PS, glycerophosphoserines; PC, glycerophosphocholines; PA, glycerophosphates; PI, glycerophosphoinositols; PGP, glycerophosphoglycerophosphates; LysoPE, lysophosphatidylethanolamines; LysoPA, lysophosphatidic acid; LysoPC, lysophosphatidylcholines; FA, fatty acids.

At 3 h the impact of the C_12_-o-(BG-D)-10 on the glycerophospholipid levels continued, as manifested by significant decreases of *sn*-glycerol 3-phosphate (log_2_FC = -2.02) and *sn*-glycero-3-phosphocholine (log_2_FC = -2.4), albeit less than at 1 h **(Fig. 1b)**. CDP-glycerol was not detected in the later time points i.e., 3 and 6 h. Choline phosphate (log_2_FC = -3.7) was further significantly reduced. Notably, abundance of glycerophospholipids and fatty acids including PG(25:0), PG(30:0), FA (17:0), FA (20:0), FA (20:1), FA methyl(14:0), FA methyl(18:0), and LysoPC(18:1) was significantly increased (≥1-log_2_FC, *p*<0.05) except for FA oxo(19:0) (≥ -1-log_2_FC, *p*<0.05). Importantly, the abundance of *N*-butyryl-L-homoserine lactone, a quorum-sensing signaling molecule, was markedly decreased (log_2_FC = -1.00) in response to C_12_-o-(BG-D)-10 treatment **(Fig. 1b)** [47].

At 6 h, the level of *sn*-glycero-3-phosphocholine decreased moderately (log_2_FC = -0.68), while *sn*-glycerol 3-phosphate was completely diminished and was not detected. The level of choline phosphate (log_2_FC = -1.8) was reduced. Most fatty acids and phosphatidylglycerol phosphate levels were further increased, including PG (30:0), FA hydroxy (4:0), FA hydroxy (8:0), LysoPC (18:1), and PGP (36:5) (≥ 1-log_2_FC, *p*<0.05). The level of (R)-3-hydroxybutanoate was further significantly decreased at 6 h (log_2_FC = -1.06). Also, the levels of [FA (21:2)] octadecenoic acid, and FA methyl (18:0) were reduced (≥ -1-log_2_FC, *p*<0.05) **(Fig. 1c)**.

Given that a higher number of significant perturbations of the lipid bilayer was observed at 1 h compared to 3 and 6 h, this suggests that C_12_-o-(BG-D)-10 disrupts the lipid bacterial membrane as the first step of the bacterial killing mechanism.

### Amino-sugar and sugar-nucleotide metabolism, peptidoglycan and teichoic acid biosynthesis

Sugar nucleotides and amino sugars are the building blocks of peptidoglycans, involved in the synthesis of bacterial cell walls and teichoic acid in Gram-positive bacteria [48]. After C_12_-o-(BG-D)-10 treatment, sugar nucleotides and amino sugar metabolism showed substantial alterations, indicating down regulation of peptidoglycan biosynthesis across all time points (1, 3 and 6 h). At 1 h, the treated bacterial cells displayed significant perturbations of metabolites involved in the early stages of peptidoglycan and teichoic acid formation. In contrast, modest disturbances were observed at 6 h **(Fig. 2a)**.

**Figure 2.**
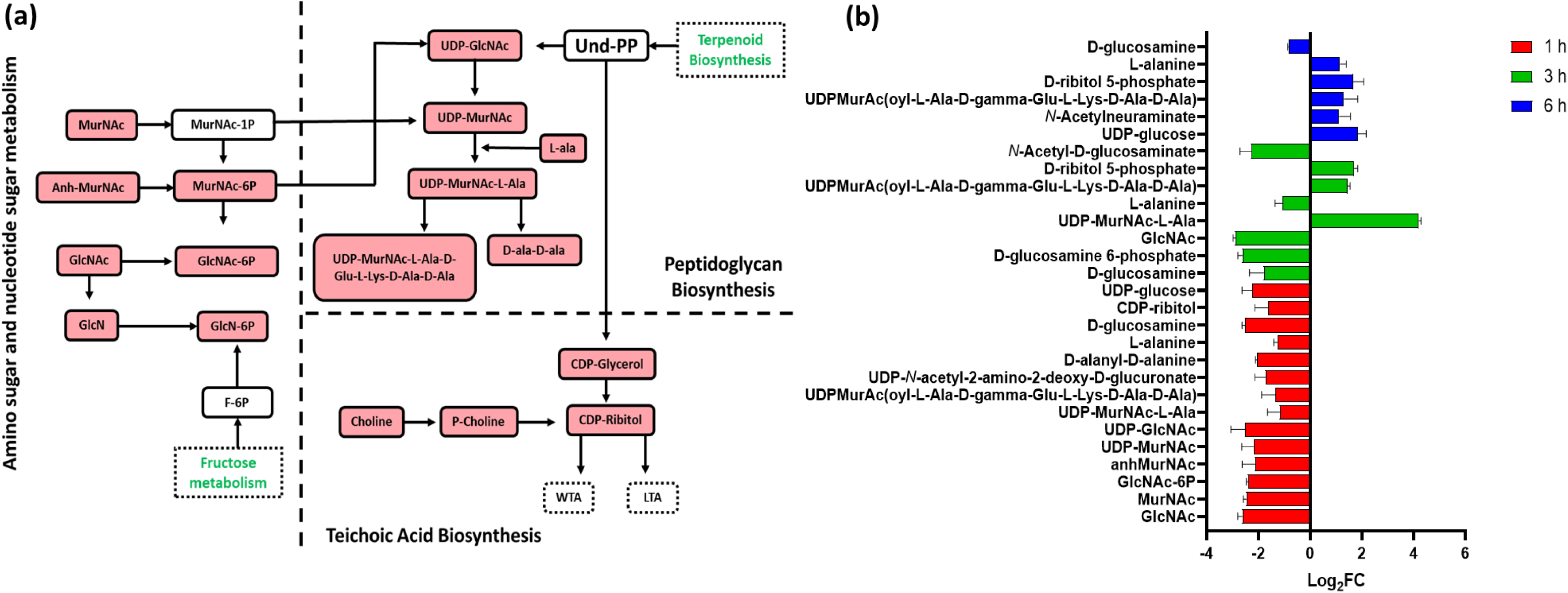
**(a)** Diagrammatic representation of all significantly impacted [increased: blue, decreased: red] amino-sugar and sugar-nucleotide metabolites in MRSA ATCC 43300 following treatment with C_12_-o-(BG-D)-10. **(b)** Significantly impacted amino-sugar and sugar-nucleotides in MRSA ATCC 43300 following treatment with C_12_-o-(BG-D)-10 at 1 h (red), 3 h (green) and 6 h (blue). Putative metabolite names are assigned based on accurate mass (≥1.0-log_2_-FC; *p*<0.05).

At 1 h after C_12_-o-(BG-D)-10 treatment, the levels of seven crucial precursors involved in the biosynthesis of bacterial cell walls exhibited a significant decline. These included *N*-acetyl-D-glucosaminate (GlcNAc), *N*-acetylmuramate (MurNAc), *N*-acetyl-D-glucosamine 6-phosphate (GlcNAc-6P), 1,6-anhydro-*N*-acetylmuramate (anhMurNAc), UDP-*N*-acetylmuramate (UDP-MurNAc) and UDP-*N*-acetyl-D-glucosamine (UDP-GlcNAc) (≥ -2-log_2_FC, *p*<0.05) **(Fig. 2b)**. Likewise, the precursors of the downstream pathway, peptidoglycan, including D-alanyl-D-alanine, UDP-*N*-acetylmuramoyl-L-alanine (UDP-MurNAc-L-Ala), UDP-MurAc(oyl-L-Ala-D-gamma-Glu-L-Lys-D-Ala-D-Ala), UDP-*N*-acetyl-2-amino-2-deoxy-D-glucuronate and L-alanine were also markedly reduced (≥ -1-log_2_-FC, *p*<0.05). Interestingly, C_12_-o-(BG-D)-10 caused a significant impact on the crucial elements of wall teichoic acid. Notably, it decreased the levels of key constituents namely MurNAc and GlcNAc, which also play integral roles in amino sugar metabolism, as well as the level of CDP-ribitol, a pivotal molecule of wall teichoic acid biogenesis (≥ -1.0-log_2_FC, *p*<0.05) [49]. The level of choline involved in the synthesis of crucial intermediates of teichoic acid was also significantly reduced (log_2_FC = -3.32) [50]. UDP-glucose, a nucleotide sugar which serves as a key intracellular intermediary in biosynthesis of the bacterial cell envelope, was significantly lowered (log_2_FC = -2.1) at 1 h **(Fig. 2b)** [51].

At 3 h, further perturbations in the building blocks of peptidoglycan were observed, where the levels of amino and nucleotide sugars intermediates involved in generation of peptidoglycan building blocks were significantly reduced. These included, D-glucosamine 6-phosphate, *N*-acetyl-D-glucosamine, and GlcNAc which were reduced (≥ -2-log_2_FC, *p*<0.05) **(Fig. 2b)**. Additionally, the level of UDP-MurNAc-L-Ala and UDP-MurNAc(oyl-L-Ala-D-gamma-Glu-L-Lys-D-Ala-D-Ala) was increased (≥ 1-log_2_FC, *p*<0.05) while the level of D-glucosamine and L-alanine was decreased (≥ -1-log_2_FC, *p*<0.05). The level of D-ribitol 5-phosphate was significantly increased (log_2_FC = 1.7), which has a crucial role in the synthesis of wall teichoic acid in the bacterial cell envelope **(Fig. 2b)** [41].

Contrary to the pronounced effects observed at earlier time points, 1 and 3 h, C_12_-o-(BG-D)-10 treatment caused a minor effect on sugar nucleotides and amino sugars and its downstream pathways (peptidoglycan and wall teichoic acid) at 6 h. Notably, there was a significant increase in the levels of UDP-glucose, N-acetylneuraminate, UDP-MurNAc(oyl-L-Ala-D-gamma-Glu-L-Lys-D-Ala-D-Ala), D-ribitol 5-phosphate, and L-alanine (≥ 1-log_2_FC, *p*<0.05) **(Fig. 2b)**.

Similar to response patterns of glycerophospholipids and fatty acids, the amino and nucleotide sugars and metabolites involved in the biogenesis of peptidoglycan were more frequently perturbed at 1 h than at later time points.

### Histidine metabolism

The histidine metabolic pathway plays a crucial role in fundamental regulatory processes in bacteria, including amino-acids, purines and thiamine biosynthesis [52]. The main metabolic intermediate generated by the histidine pathway is 5′-phosphoribosyl-4-carboxamide-5-aminoimidazole (AICAR). This intermediate is at the crossroads between purine-histidine cross-talk [53]. Therefore, the histidine pathway has been extensively investigated as a promising therapeutic target for novel antibiotics to treat infections caused by Staphylococcus [54, 55]. Treatment with C_12_-o-(BG-D)-10 led to perturbation in several essential metabolites within the histidine biosynthetic pathway, notably including intermediates like L-glutamine and L-glutamate, both recognized as indicators of bacterial stress response **(Fig. 3a)** [56].

**Figure 3.**
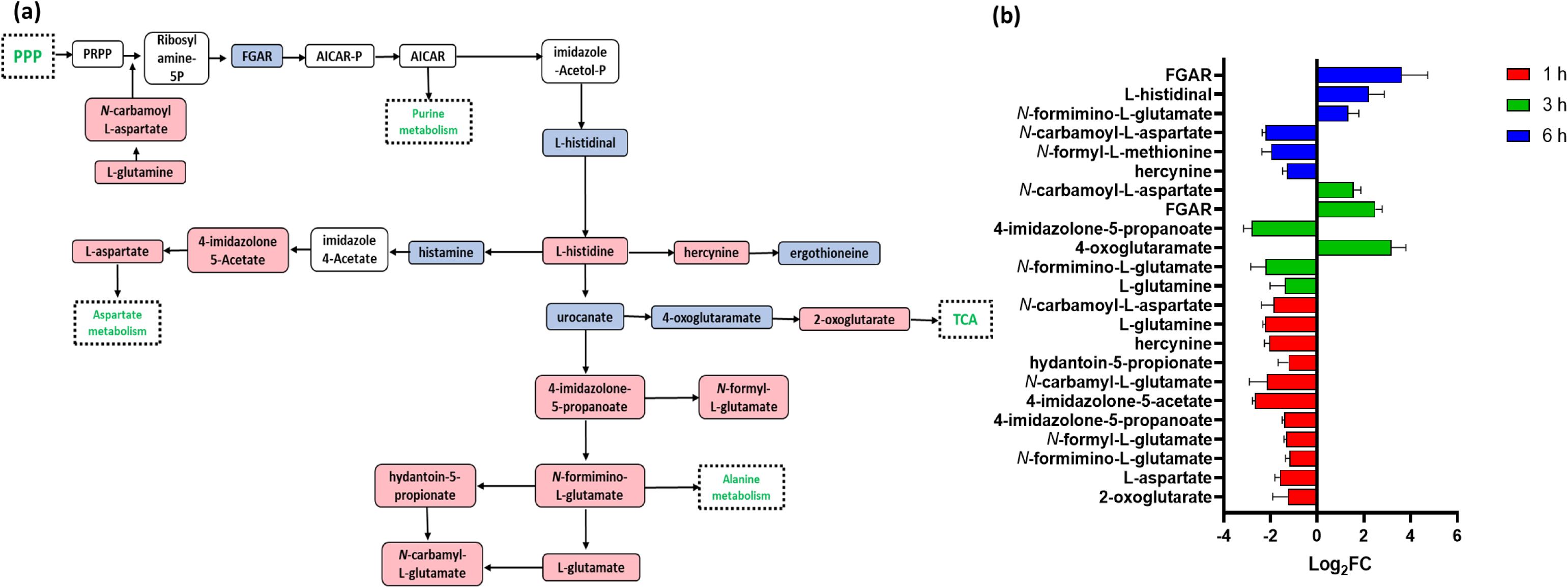
**(a)** Diagrammatic representation of all significantly impacted [increased: blue, decreased: red] histidine metabolites in MRSA ATCC 43300 following treatment with C_12_-o-(BG-D)-10. **(b)** Significantly impacted histidine metabolites in MRSA ATCC 43300 following treatment with C_12_-o-(BG-D)-10 at 1 h (red), 3 h (green), and 6 h (blue). Putative metabolite names are assigned based on accurate mass (≥ 1.0-log_2_-FC; *p*<0.05).

At 1 h post C_12_-o-(BG-D)-10 treatment, ten essential intermediates of histidine metabolism were significantly perturbed. Specifically, 4-imidazolone-5-acetate, *N*-carbamyl-L-glutamate, hydantoin-5-propionate, L-glutamine and hercynine displayed more significant alteration (≥ - 2-log_2_FC, *p*<0.05) compared to 4-imidazolone-5-propanoate, *N*-Formyl-L-glutamate, *N*-formimino-L-glutamate, L-aspartate and 2-oxoglutarate (≥ -1-log_2_FC, *p*<0.05) **(Fig. 3b)**.

At 3 h, alterations were more pronounced than at 1 h. This was evident in decreased abundance of L-glutamate, L-glutamine, *N*-formimino L-glutamate and 4-imidazolone-5-propanoate (≥ - 1-log_2_FC, *p*<0.05), accompanied by an increased level of 4-oxoglutaramate (log_2_FC = 3.1). Importantly, the level of 5-phosphoribosyl-*N*-formylglycinamide (FGAR) (log_2_FC = 2.4) significantly increased, a key element in AICAR synthesis **(Fig. 3b)**.

At 6 h, further significant perturbations were observed in histidine biosynthetic pathway. FGAR, N-formimino-L-glutamate and L-histidinal levels increased (≥ 1-log_2_FC, *p*<0.05), while the levels of hercynine, *N*-formyl-L-methionine, and *N*-carbamoyl-L-aspartat decreased (≥ -1-log_2_FC, *p*<0.05) **(Fig. 3b)**. These time-sensitive changes underscore a dynamic response of the histidine pathway to the treatment.

### Nucleotide (purine and pyrimidine) metabolism

C_12_-o-(BG-D)-10 treatment induced a marked dysregulation in the histidine interconnected pathway, nucleotide (purine and pyrimidine) metabolism, both are crucial for DNA and RNA formation **(Fig. 4a)** [57]. This impact was more pronounced at 1 and 3 h compared to the later time point, 6 h **(Fig. 4b)**.

**Figure 4.**
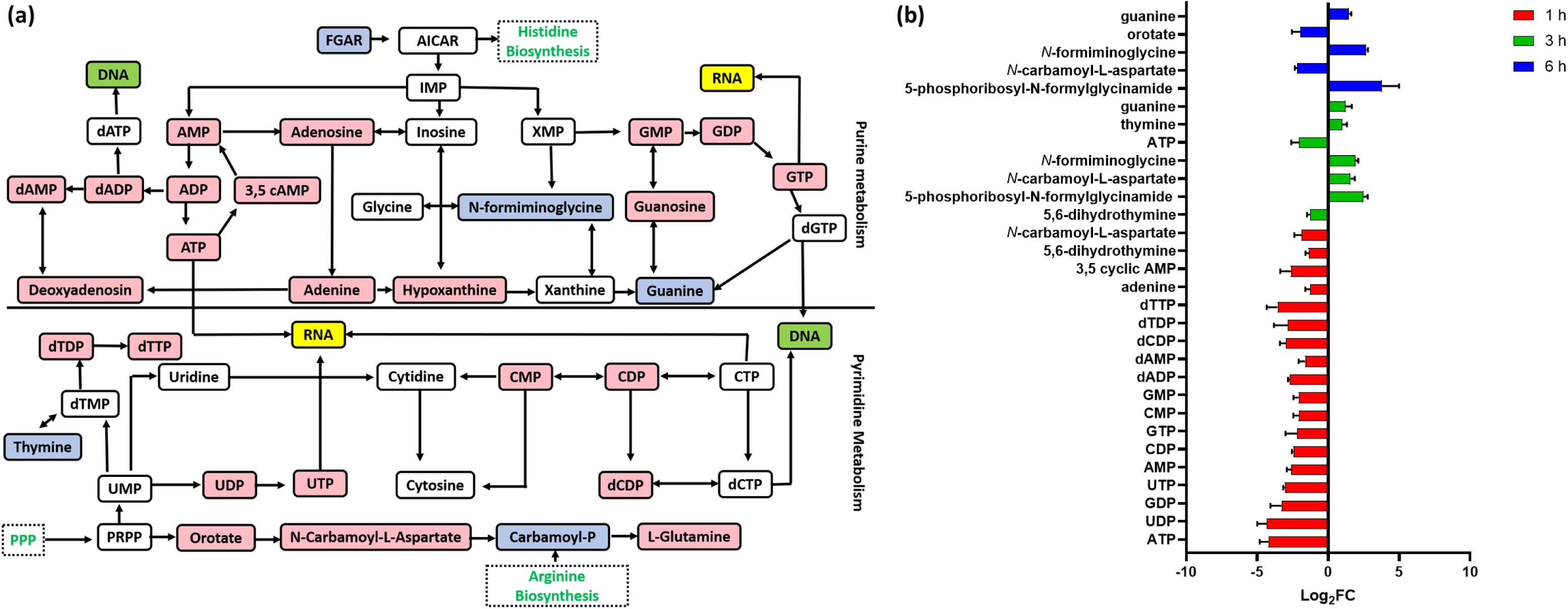
**(a)** Diagrammatic representation of all significantly impacted [increased: blue, decreased: red] pyrimidine and purine metabolites in MRSA ATCC 43300 following treatment with C_12_-o-(BG-D)-10. **(b)** Significantly impacted nucleotide and peptide metabolites in MRSA ATCC 43300 following treatment with C_12_-o-(BG-D)-10 at 1 h (red), 3 h (green), and 6 h (blue). Putative metabolite names are assigned based on accurate mass (≥1.0-log_2_-FC; *p*<0.05).

At 1 h, a marked decline in the levels of 14 nucleotides was observed following C_12_-o-(BG-D)-10 treatment, including ATP, UTP, AMP and dAMP (≥ -1.5-log_2_FC, *p*<0.05), which ultimately impacted DNA and RNA synthesis **(Fig. 4b)** [58, 59]. Among nucleotide bases, adenine (log_2_FC = -1.2) was significantly reduced **(Fig. 4b)**. An important observation at 1 h was that the level of 3,5 cyclic AMP required for cyclic di-AMP synthesis was profoundly reduced (log_2_FC = -2.7) **(Fig. 4b)**. Cyclic di-AMP is an essential regulator for several bacterial biochemical processes, including metabolic pathways governing bacterial growth (fatty acid, carbohydrates and nucleotide metabolism), virulence, biofilm formation and cell cycle progression [60]. It is also involved in bacterial cell survival mechanism [61]. The level of 5,6-dihydrothymine was reduced (log_2_FC = -1.6). Furthermore, the level of *N*-carbamoyl-L-aspartate another key intermediate in the metabolism of purines and pyrimidines was also decreased (log_2_FC = -1.91) **(Fig. 4b)** [62].

At 3 h, significant perturbations were observed in the levels of two pivotal intermediates of purines and pyrimidines metabolism. Specifically, the level of 5,6-dihydrothymine (log_2_FC = -1.2) exhibited a decrease, while the level of 5-phosphoribosyl-*N*-formylglycinamide (log_2_FC = 2.4) showed an increase [62, 63]. Moreover, the levels of *N*-carbamoyl-L-aspartate and *N*-formiminoglycine, thymine and guanine demonstrated increases (≥ 1.0-log_2_FC, *p*<0.05) **(Fig. 4b)**. However, the level of ATP, which is the major source of energy in the bacteria, experienced a further significant reduction (log_2_FC = -1.9) [64].

At 6 h, a notable increase was observed in the level of 5’-phosphoribosyl-*N*-formylglycinamide (log_2_FC = 3.7). Conversely, the *N*-carbamoyl-L-aspartate level experienced a further decline (log_2_FC = -2.2), while the level of *N*-formiminoglycine increased (log_2_FC =2.6) **(Fig. 4b)**. Orotate (log_2_FC = -1.9), an essential metabolite in the biosynthesis of pyrimidine, was significantly decreased at 6 h **(Fig. 4b)** [65]. Among nucleotide bases, the level of guanine (log_2_FC = 1.4) was significantly increased **(Fig. 4b)**.

### Central carbon metabolism

Central carbon metabolism, which includes glycolysis, pentose phosphate pathway (PPP) and tricarboxylic acid cycle (TCA), is required for bacteria to generate energy in the form of ATP. This process provides precursors for all the biosynthetic reactions that are required for cell survival [66, 67]. This makes central carbon metabolism i.e., carbon uptake and carbon utilization, an attractive antimicrobial target because these processes are typically essential for microbial survival [68–70]. Innate immunity, which frequently deprives bacteria of iron, amino acids, and other essential nutrients, serves as an example of the efficacy of this strategy [71–73]. C_12_-o-(BG-D)-10 treatment caused perturbations in all three pathways of central carbon metabolism more significant at 1 h compared to later timepoints **(Fig. 5a)**.

**Figure 5.**
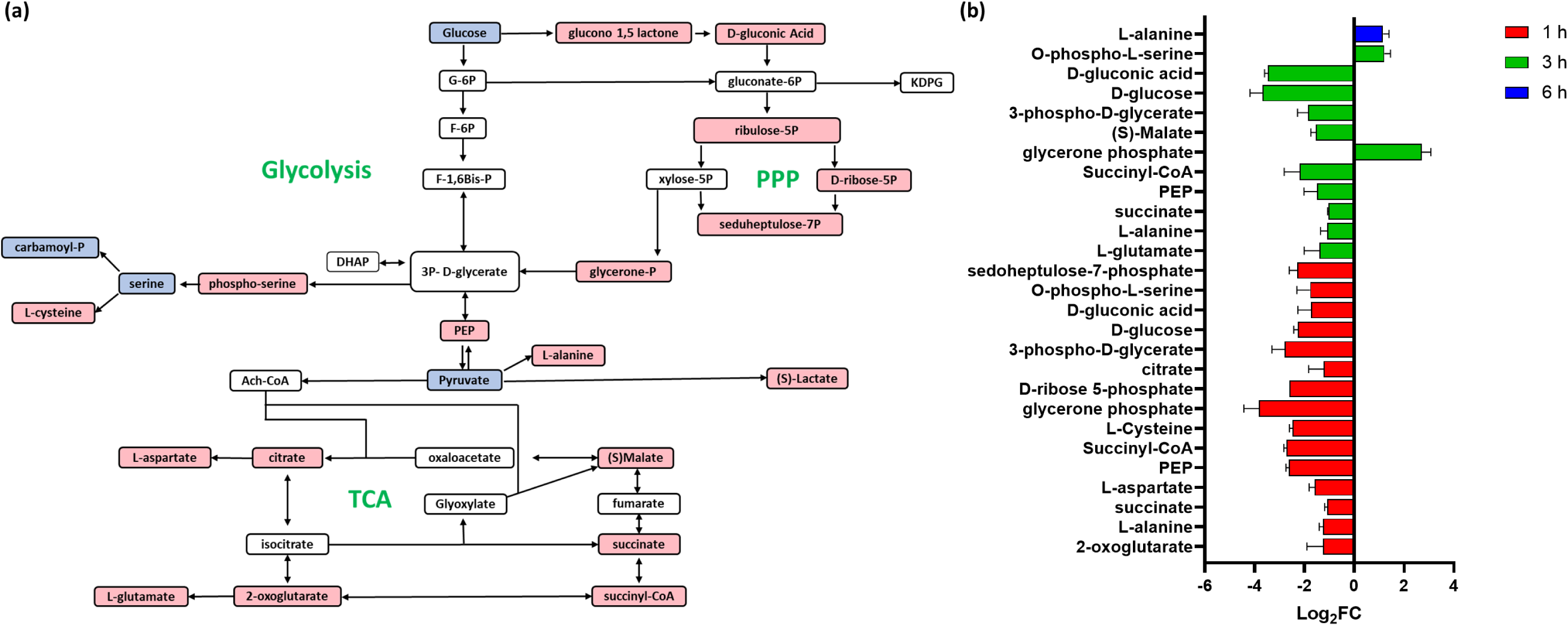
**(a)** Diagrammatic representation of all significantly impacted [increased: blue, decreased: red] central carbon metabolism metabolites in MRSA ATCC 43300 following treatment with C_12_-o-(BG-D)-10. **(b)** Significantly impacted central carbon metabolism metabolites in MRSA ATCC 43300 following treatment with C_12_-o-(BG-D)-10 at 1 h (red), 3 h (green), and 6 h (blue). Putative metabolite names are assigned based on accurate mass (≥1.0-log_2_-FC; *p*<0.05).

At 1 h, 18 important intermediates of central carbon metabolism, interlinking glycolysis, TCA and PPP were significantly perturbed **(Fig. 5b)**. This included, glycerone phosphate, phosphoenolpyruvate (PEP), succinyl-CoA, D-ribose 5-phosphate, 3-phospho-D-glycerate, L-cysteine, D-glucose, and sedoheptulose-7-phosphate, 2-oxoglutarate, L-alanine, succinate, L-aspartate, citrate, D-gluconic acid, and O-phospho-L-serine (≥ -1-log_2_FC, *p*<0.05) **(Fig. 5b)**.

The effect of C_12_-o-(BG-D)-10 continued at 3 h, when D-glucose, D-gluconic acid, L-glutamate, L-alanine, succinate, PEP, (S)-Malate, 3-phospho-D-glycerate and Succinyl-CoA (≥ -1-log_2_FC, *p*<0.05) were significantly perturbed. O-phospho-L-serine and glycerone phosphate (≥ 1-log_2_FC, *p*<0.05) and were also perturbed **(Fig. 5b).**

At 6 h, most of the metabolites of the central carbon metabolism were diminished and were not detected except for L-alanine which was further significantly perturbed (log_2_FC = 1.15) **(Fig. 5b).**

Taken together, the crucial central carbon metabolism was largely and significantly perturbed at 1 h and 3 h while only minor perturbations were observed at 6 h.

### Arginine metabolism

The disruption of arginine (one of the most versatile and inter-convertible) metabolism has recently emerged as a powerful approach to control and subvert bacterial pathogenesis [74]. C_12_-o-(BG-D)-10 treatment significantly impacted arginine metabolism, particularly at 1 and 3 h, with minimal changes observed after 6 h of treatment. **(Fig. 6a)**.

**Figure 6.**
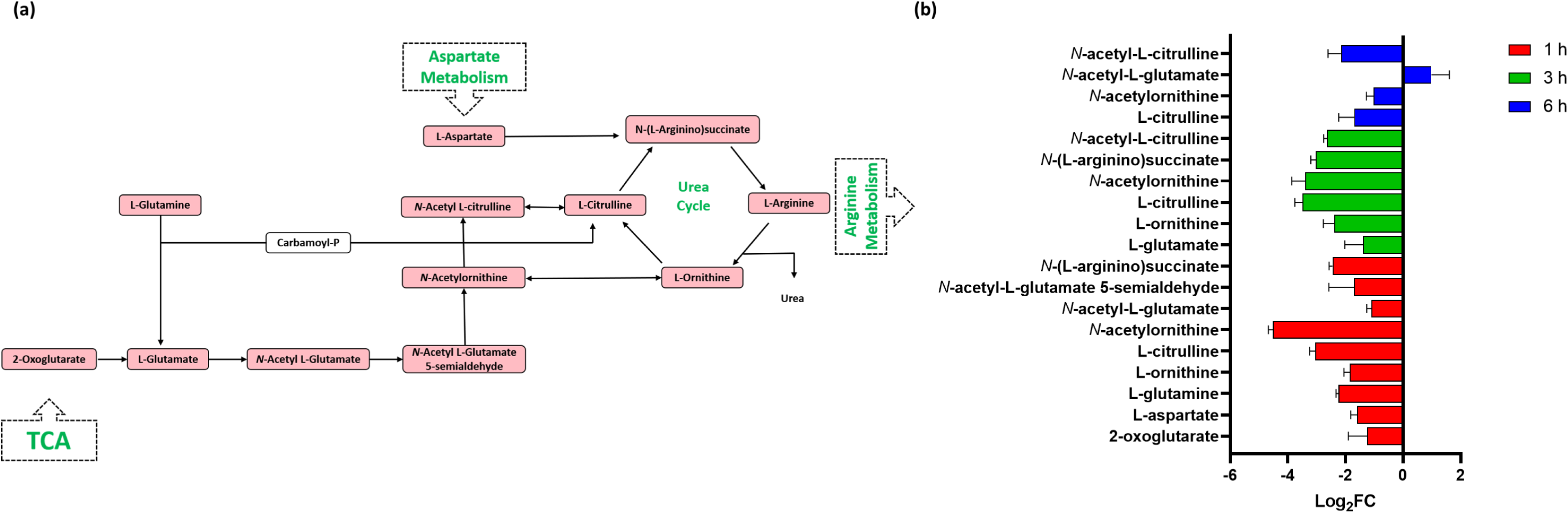
**(a)** Diagrammatic representation of all significantly impacted [increased: blue, decreased: red] arginine metabolism and interrelated TCA cycle metabolites in MRSA ATCC 43300 following treatment with C_12_-o-(BG-D)-10. **(b)** Significantly impacted metabolites in the arginine biosynthesis and metabolism in MRSA ATCC 43300 following treatment with C_12_-o-(BG-D)-10 at 1 h (red), 3 h (green), and 6 h (blue). Putative metabolite names are assigned based on accurate mass (≥1.0-log_2_-FC; *p*<0.05).

After 1 h of treatment, nine metabolites of arginine biosynthesis, including L-aspartate, L-glutamine, L-citrulline, L-ornithine, *N*-acetyl-L-glutamate, *N*-(L-arginino) succinate, *N*-acetyl-L-glutamate 5-semialdehyde, 2-oxoglutarate and *N*-acetylornithine, were significantly perturbed (≥ -1-log_2_-FC, *p*<0.05) **(Fig. 6b)**.

At 3 h, the inhibitory impact of C_12_-o-(BG-D)-10 on arginine biosynthesis continued, although at intensity wherein the levels of six important intermediates, including L-glutamine, L-ornithine, L-citrulline, *N*-(L-arginino) succinate, *N*-acetyl-L-citrulline and *N*-acetylornithine were significantly reduced (≥ -1-log_2_-FC, *p*<0.05) **(Fig. 6b)**.

By 6 h, the level of *N*-acetyl-L-glutamate was increased (log_2_FC =1.0), signifying a bacterial stress response. The levels of L-citrulline, *N*-acetylornithine and *N*-acetyl-L-citrulline were decreased (≥ -1-log_2_FC, *p*<0.05) **(Fig. 6b)**.

### Coenzyme A (CoA) biosynthesis

Pantothenic acid, also known as vit-B5, is essentially required by bacteria to synthesize CoA. CoA, in turn, is crucial for generation of fatty acids, carbohydrates, proteins, and even some intermediates within the TCA cycle [75]. C_12_-o-(BG-D)-10 treatment resulted in marked disruption in the pantothenate and coenzyme A (CoA) biosynthesis **(Fig. S7a)**.

At 1 h, C_12_-o-(BG-D)-10 treatment caused significant perturbations in the levels of five metabolites related to the pantothenate and CoA biosynthesis pathways **(Fig. S7b)**. In particular, the levels of L-cysteine, CoA, L-aspartate and 5,6-dihydrouracil significantly decreased (≥ -1.0-log_2_FC, *p*<0.05) **(Fig. S7b)**. In contrast, uracil level increased (log_2_FC =1.04). At 3 h, there was a notable decrease (≥ -2-log2FC, p<0.05) in the *N*-((R)-pantothenoyl)-L-cysteine and (R)-2,3-dihydroxy-3-methylbutanoate. Additionally, the level of pantothenate decreased (log_2_FC = -1.5) whereas 3-methyl-2-oxobutanoic acid increased (log_2_FC = 1.4) **(Fig. S7b)**. The effect of C_12_-o-(BG-D)-10 persisted at 6 h, when the levels of pantetheine and pantothenate increased (≥ 1-log_2_FC, *p*<0.05), while *N*-((R)-pantothenoyl)-L-cysteine demonstrated decreases (log_2_FC = -2.6) **(Fig. S7b)**.

### Homeostasis and stress metabolism

C_12_-o-(BG-D)-10 treatment induced a significant perturbation in the metabolites responsible for bacterial homeostasis. D-glucose-6-phosphate, a substrate for trehalose-6-phosphate and UDP-glucose synthesis, which experienced marked perturbations across all timepoints: at 1 h (log_2_FC = -1.8), 3 h (log_2_FC = -0.7) and 6 h (log_2_FC = 1.1) **(Fig. S8)** [76, 77]. Trehalose, in its dephosphorylated form, serves as a recognized osmoprotectant in bacteria during osmotic stress [78]. Significant alteration in the level of trehalose-6-phosphate at 3 h (log_2_FC = -1.71) indicated that the C_12_-o-(BG-D)-10 treatment induced osmotic stress. Choline, a precursor to glycine betaine (GB), which acts as a potent osmoprotectant underwent significant reduction at 1 h (log_2_FC = -3.3) [79]. However, choline was not detected at the later timepoints. Likewise, PPP-intermediates involved in the maintenance of redox homeostasis in the cell were markedly perturbed, which was indicated by a significant decrease in the levels of ribose-5-phosphate, NADP+ and CMP (≥ -1.5-log_2_FC, *p*<0.05) at 1 h **(Fig. S8)** [80]. However, these PPP intermediates were not detected at later timepoints.

## Conclusion

MRSA is a serious threat to global heath, and contributes to nearly half a million annual deaths. As further resistance emerges against current antimicrobials in clinical use, there is an urgent need for new treatment options. Our group has synthesized CLOs modeled on the structure of antimicrobial peptides. C_12_-o-(BG-D)-10 was previously found to inhibit the growth of MRSA *in vitro*, however this activity was only partially attributable to membrane disruption as evident *via* fluorescence assay. The full mechanism(s) of its antimicrobial activity was not fully understood. This is the first study to investigate the mechanism(s) of antimicrobial action of a CLO i.e., C_12_-o-(BG-D)-10 using metabolomics. C_12_-o-(BG-D)-10 antimicrobial action commences with disrupting the bacterial cell envelope. Early at 1 h, significant perturbations were observed in cell membrane lipids and glycerophospholipids, along with sugar nucleotides and amino-sugar metabolites linked to peptidoglycan and teichoic acid biosynthesis. The polymer also affected RNA and DNA biosynthesis and led to pronounced perturbations in histidine metabolism (linked to the synthesis of purines and pyrimidines), energy metabolism (i.e. arginine and TCA cycle), pantothenate biosynthesis and CoA biogenesis (essentially required by cells for survival and normal growth). C_12_-o-(BG-D)-10 also perturbed central carbon metabolism and the stress pathway in the bacteria more prominently at the initial time points i.e., 1 and 3 h. These insights on the mechanisms of action of C_12_-o-(BG-D)-10 will enable the rational design of antimicrobial combinations of clinically available antimicrobials with C_12_-o-(BG-D)-10 in future *in vitro* and *in vivo* studies in an approach to achieve synergistic and effective bacterial killing.

## MATERIALS AND METHODS

### CLO and antibiotic stock solution, media and bacterial isolates

C_12_-o-(BG-D)-10 was synthesized by Cu(0)-mediated reversible deactivation radical polymerization (RDRP) using the protocol by Grace *et al*. [29]. The CLO stock solutions were prepared by dissolving the CLO in DMSO first, diluting with MilliQ water (to 20% DMSO) and then vortexing until clear. DMSO was filtered through 0.22µm sterile nylon filters before use. Methicillin-resistant *S. aureus* (MRSA) ATCC 43300 was used in the study. All susceptibility and time–kill studies were performed in cation-adjusted Mueller-Hinton broth (CAMHB; containing 20-25 mg/L Ca^2+^ and 10-12.5 mg/L Mg^2+^; BD, Sparks, MD, USA). Viable counting was performed on cation-adjusted Mueller–Hinton agar (CAMHA; containing 25 mg/L Ca^2+^ and 12.5 mg/L Mg^2+^; BD, Sparks, MD, USA).

### Antibacterial killing kinetics of CLO

A static concentration time-kill assay of C_12_-o-(BG-D)-10 against MRSA ATCC 43300 was performed **(Fig. S2)**. Before the time-kill assay, MRSA ATCC 43300 was sub-cultured on a CAMHA plate and then incubated at 37°C for ∼18-24 h. Three colonies were transferred from the CAMHA plate to inoculate 10 mL of sterile CAMHB in a 50 mL Falcon tube and incubated overnight in a shaking water bath (37°C, 150 rpm, ∼16 h). The optical density of the bacterial suspension was measured using a spectrophotometer, and the suspension was appropriately diluted to achieve the targeted initial inoculum of ∼10^6^ CFU/mL **(Fig. S2b)**. The inoculated tubes were dosed with C_12_-o-(BG-D)-10 (in 20% DMSO) to achieve concentrations of 8, 16 and 64 µg/mL. The DMSO concentrations in the tubes were ≤0.125%. One culture tube was drug-free to represent the control. At 0, 1.5, 5, 24, 48 and 72 h, 1-mL samples were collected from each tube, centrifuged and washed twice with 0.9% normal saline. The samples were then serially diluted in saline plates and plated onto CAMHA plates. After 24 h incubation at 37°C, the CFU were counted, and the time-kill curves graphed as log_10_ CFU/mL vs time (hours).

### Metabolomics sample preparation

An untargeted metabolomics study was carried out to explore the mechanism(s) of action of C_12_-o-(BG-D)-10 against MRSA ATCC 43300 using a concentration of 48 µg/mL (i.e. 3×MIC). Samples were taken and analyzed at the 1-, 3-, and 6-h time points in 4 biological replicates. An overnight culture was prepared by inoculating a single colony into 100 mL CAMHB in 250 mL conical flasks (Pyrex) and incubating the suspension in a shaker at 37°C and 180 rpm for ∼16 h. After incubation overnight, log-phase cells were prepared in fresh MHB and then incubated for 2 h at 37°C at 180 rpm to log phase with a starting bacterial inoculum of 10^8^ CFU/mL. Then, C_12_-o-(BG-D)-10 was added to obtain the desired concentration of 48 µg/mL (3 x MIC), in parallel to a CLO-free control for each replicate. The flasks were then incubated at 37°C with shaking at 180 rpm. At each time point (0, 1, 3, and 6 h), 15-mL samples were transferred to 50-mL Falcon tubes for quenching, and the optical density reading at 600 nm (OD600) was then measured and normalized to the pre-treatment level of approximately ∼0.5 with fresh CAMHB. Samples were then centrifuged at 3,220 *g* and 4°C for 10 min, and the supernatants were removed. The pellets were stored at -80°C until metabolite extraction. The experiment was performed in 4 biological replicates to reduce the bias from inherent random variation.

### Metabolomics metabolite extraction

The bacterial pellets were washed twice in 1 mL of 0.9% saline and then centrifuged at 3,220 *g* and 4°C for 5 min to remove residual extracellular metabolites and medium components. The washed pellets were resuspended in a cold extraction solvent (chloroform-methanol-water at 1:3:1, vol/vol) containing 1 µM each of the internal standards 3-[(3-cholamidopropyl)-dimethylammonio]-1-propanesulfonate (CHAPS), *N*-cyclohexyl-3-aminopropanesulfonic acid (CAPS), piperazine-*N*, *N*-bis (2-ethanesulfonic acid) (PIPES), and Tris. The samples were then frozen in liquid nitrogen, thawed on ice, and vortexed to release the intracellular metabolites (3 times). Next, the samples were transferred to 1.5-mL Eppendorf tubes and centrifuged at 14,000 *g* at 4°C for 10 min to remove any particulate matter. Finally, 200 µL of the supernatant was transferred into injection vials for liquid chromatography-mass spectrometry (LC-MS) analysis. An equal volume of each sample was combined and used as a quality control (QC) sample

### LC-MS analysis of metabolites

Both hydrophilic interaction liquid chromatography (HILIC) and reversed-phase liquid chromatography (RPLC) coupled to high-resolution mass spectrometry (HRMS) were employed to ensure the detection of both hydrophilic and hydrophobic metabolites. Samples were analyzed on a Dionex U3000 high-performance liquid chromatography system (HPLC) in tandem with a Q-Exactive Orbitrap mass spectrometer (Thermo Fisher) in both positive and negative ion modes with a resolution at 35,000. The HILIC method was described previously in detail [81]. Briefly, samples maintained at 4°C were eluted through a ZIC-pHILIC column (5 μm, polymeric, 150 by 4.6 mm; SeQuant, Merck) by mobile phase A (20 mM ammonium carbonate) and mobile phase B (acetonitrile). The gradient started with 80% mobile phase B at a flow rate of 0.3 mL/min and was followed by a linear gradient to 50% mobile phase B over 15 min. The Ascentis Express C8 column (5 cm by 2.1 mm, 2.7 μm) (catalog no. 53831-U; Sigma-Aldrich) was applied in the RPLC method. The samples were controlled at 4°C and eluted by mobile phase A (40% of isopropanol and 60% of Milli-Q water with 8 mM ammonium formate and 2 mM formic acid) and mobile phase B (98% of isopropanol and 2% of Milli-Q water with 8 mM ammonium formate and 2 mM formic acid). The linear gradient started from 100% mobile phase A to a final composition of 35% mobile phase A and 65% mobile phase B over 24 min at 0.2 mL/min. All samples were analyzed within a single LC-MS batch to avoid variations. The pooled quality control samples (QC), internal standards, and total ion chromatograms were assessed to evaluate the chromatographic peaks, signal reproducibility, and stability of analytes. To assist the identification of metabolites, a mixture of ∼500 metabolite standards was analyzed within the same batch.

### Data processing, bioinformatics, and statistical analyses

IDEOM (Identification and Evaluation of Metabolites) was used to convert raw data obtained by LC-MS to annotated and hyperlinked metabolites [82]. ProteoWizard, a freely available software library for LC-MS data analysis, was first used to extract mzXML files from LC-MS raw sheets. These files were then processed using XCMS, a graphical user interface, for peak picking and generating peakML files [83]. Using MZmatch.R tool, peaks were aligned and filtered with MDI (minimum detectible intensity) of 100000 and RSD (relative standard deviation) of <0.5 and peak shape (coda dw) of >0.8. Using the same MZmatch, the missing peaks were also retrieved and annotated. Common sources of noise (contaminant signals, peak shoulders, and irreproducible peaks) were removed using the IDEOM interface. The gain and loss of protons were corrected in positive and negative electrospray ionization (ESI) mode and then a data-dependent mass recalibration (2 ppm) step for putative metabolites was performed.

Metabolites confirmed with authentic standards were assigned with MSI level 1 identification. Putative metabolites (with MSI level 2 identification) were identified by comparing their accurate masses and retention times with the standards in the databases including KEGG (Kyoto Encyclopedia of Genes and Genomics), LipidMaps, MetaCyc, and preferably EcoCyc. Peak height intensities were used for the quantification of the metabolites. Statistical analysis was performed using MetaboAnalyst 5.0 [84], a freely available online statistical tool. Briefly, putative metabolites (with median RSD ≤20% and confidence interval ≥5) were extracted from IDEOM and tabled per timepoint, then uploaded on MetaboAnalyst 5.0. Parameters were set by checking IQR (interquartile range), normalization by the median, log_2_ transformation, and autoscaling. Fold-change was calculated relative to the control from the corresponding time point. Univariate analysis was performed using two-sample t-tests [≥1 log_2_FC; t-tests, FDR adjusted *p*-value ≤0.05] to determine significantly perturbed metabolites for each timepoint. Multivariate analysis was performed and included generation of heat map and PCA plots. Lastly, the KEGG IDs of metabolites were uploaded to KEGG Mapper [85], and pathways were constructed. The individual value plots for the significantly perturbed metabolites after treatment with C_12_-o-(BG-D)-10, across all the time points can be found in the supplementary information (SI) section **(Fig. S9-S14)**.

## DATA AVAILABILITY

The entirety of our raw data has been uploaded to Metabolomics Workbench under Study No. ST003053.

## ACKNOWLEGMENTS

M.H. and M.B.H.M. both contributed equally to the preparation of this manuscript. Author order was determined both alphabetically and on the basis of seniority. This work was supported by the Australian Research Council (DP200102829), the Monash Institute of Pharmacy and Pharmaceutical Sciences (MIPS), and the National Health and Medical Research Council (GNT1159579). J.F.Q is grateful for an ARC Future Fellowship (FT170100144).

