## Supplemental materials for "Providing insight into the mechanism of action of Cationic Lipidated Oligomers (CLOs) using metabolomics"

**Supplementary Information:**

**Table S1:** Pathway enrichment analysis (performed with KEGG Mapper) of significantly perturbed metabolites across the entire study duration.

| **Pathways** | **No. of significantly perturbed metabolites** |
|---|---|
| Glycerophospholipids and FA metabolism | 20 |
| Peptidoglycan and teichoic acid biosynthesis | 18 |
| DNA and RNA biosynthesis/nucleotide biosynthesis | 23 |
| Central carbon metabolism | 19 |
| Arginine biosynthesis | 11 |
| Histidine metabolism | 15 |
| Pantothenate and CoA biosynthesis | 10 |

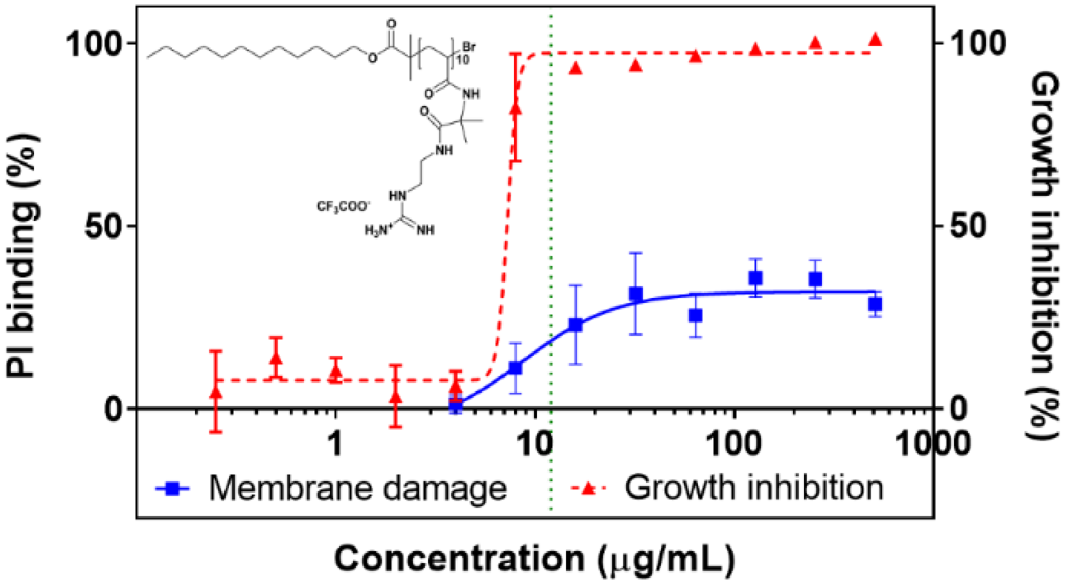


**Figure S1**. Membrane permeabilization (%; via PI binding assay) and growth inhibition (%; broth microdilution assay) of $C_{12}$-o-(BG-D)-10 versus MRSA ATCC 43300, as previously reported.

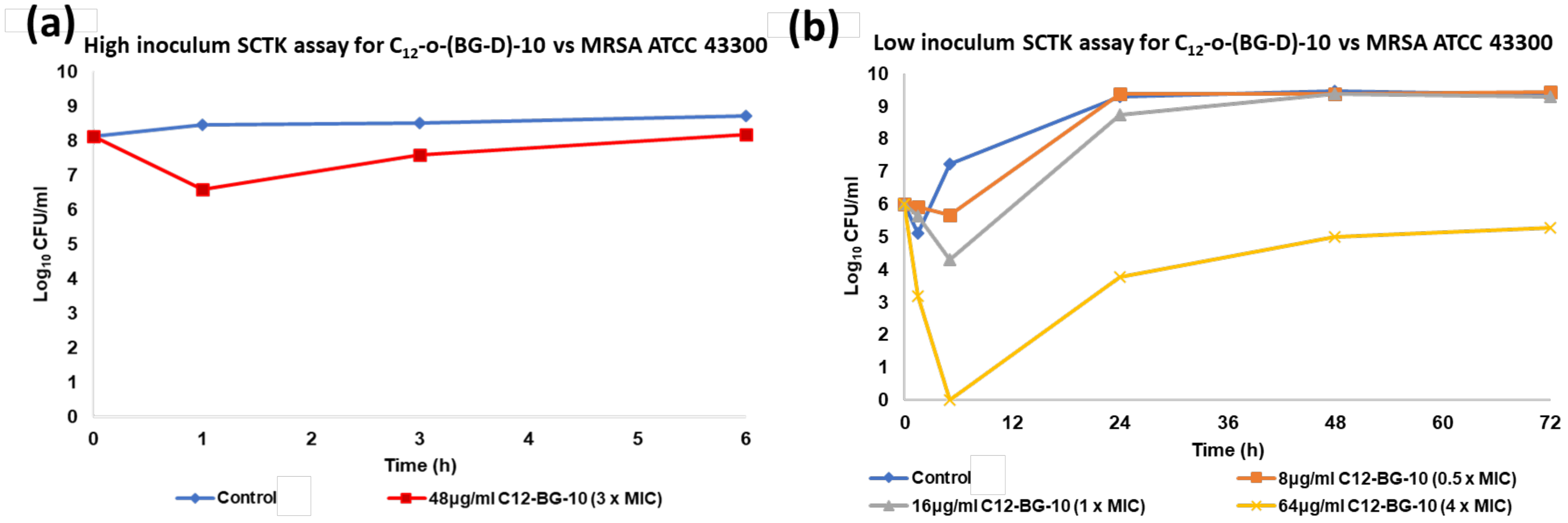


**Figure S2**. **(a)** Killing kinetics achieved in metabolomics study with 48µg/mL (3 x MIC) of $C_{12}$-o-(BG-D)-10 (pre-normalization). **(b)** Killing kinetics of MRSA ATCC 43300 (initial inoculum$10^6$ $log_{10}$ CFU/mL), after treatment with different concentrations [8µg/mL (0.5 x MIC), 16µg/mL (1 x MIC), 64µg/mL (4 x MIC)] of $C_{12}$-o-(BG-D)-10.

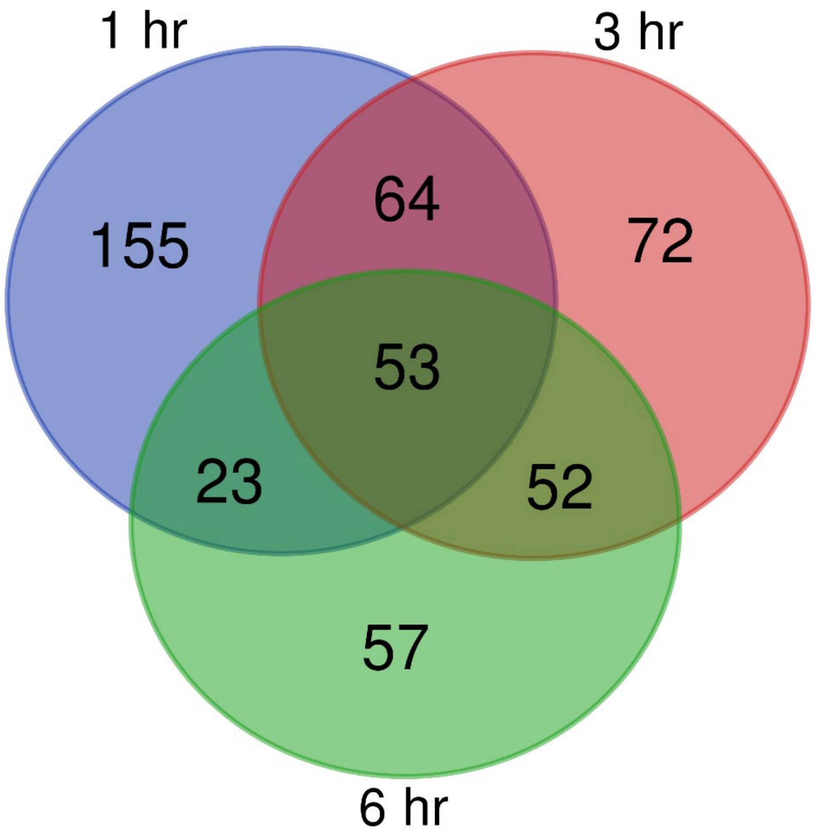


**Figure S3.** Venn diagram showing the number of metabolites of MRSA ATCC 43300, significantly affected by treatment with $C_{12}$-o-(BG-D)-10. Significant metabolites were selected with (≥ 0.5-$log_2$-FC; $P$<0.05).

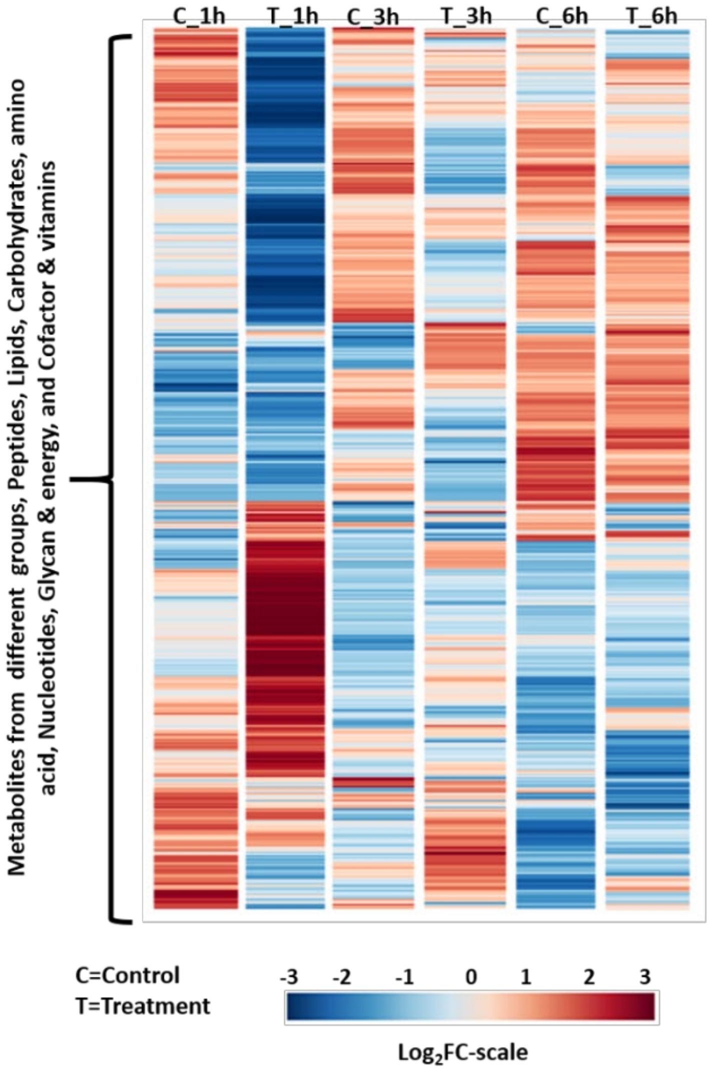


**Figure S4.** Heatmaps by treatment with 48µg/mL (3xMIC) of $C_{12}$-o-(BG-D)-10 at 1, 3 and 6 h (T_1h, T_3h and T_6h, respectively) and control (C_1h, C_3h and C_6h) in strain MRSA ATCC 43300 using untargeted metabolomics.

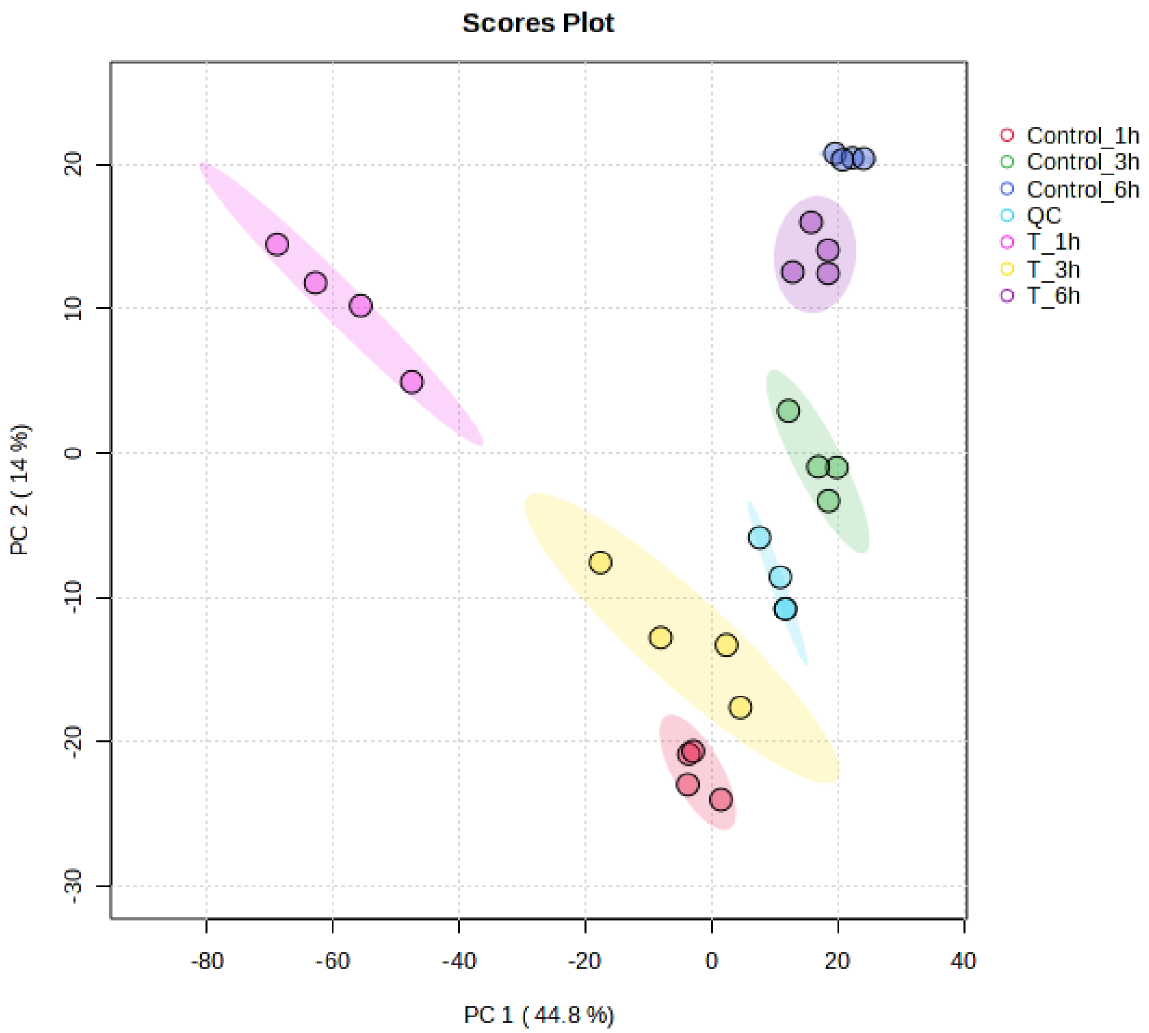


**Figure S5**. PCA plot of $C_{12}$-o-(BG-D)-10 at 1, 3 and 6 h. T= $C_{12}$-o-(BG-D)-10

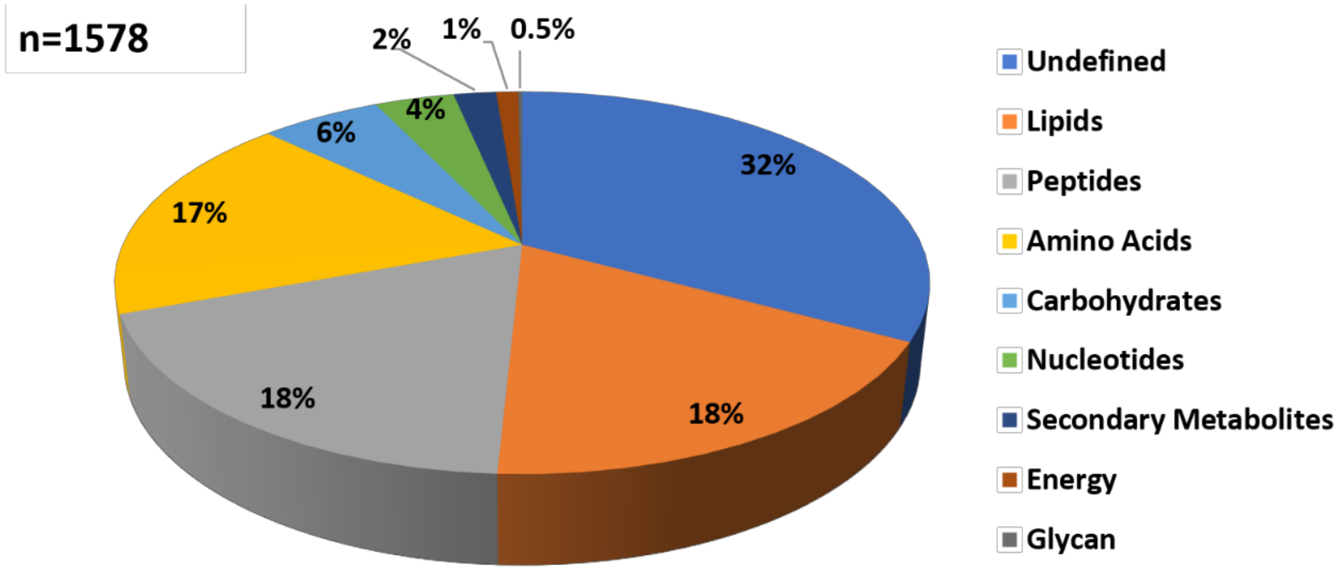


**Figure S6.** Classification of perturbed metabolites after treatment with C12-o-(BG-D)-10.

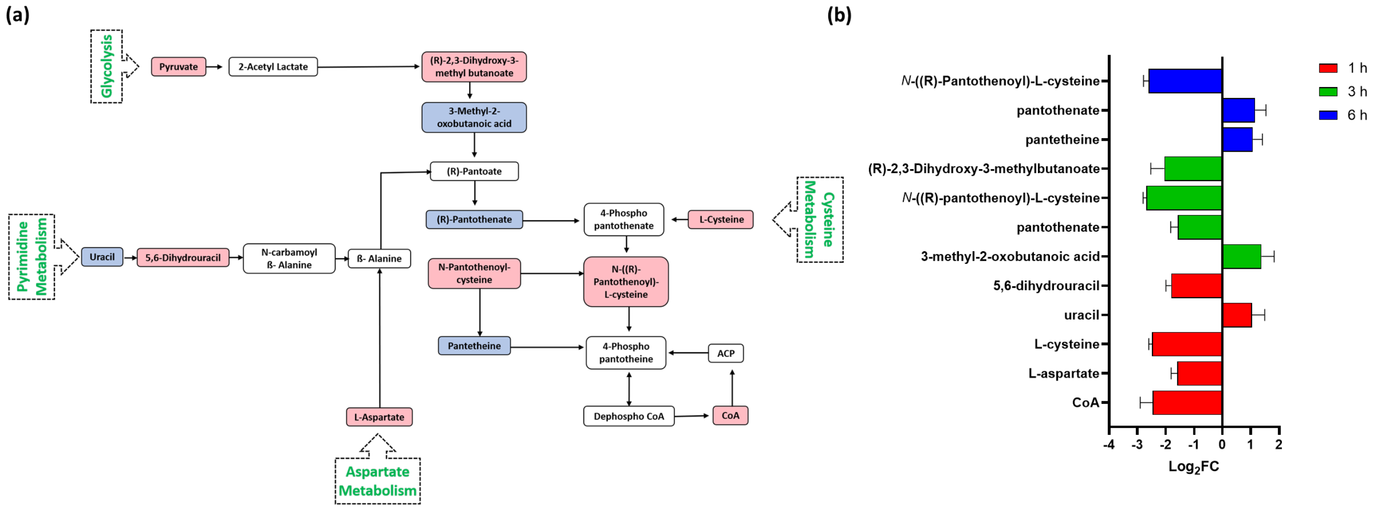


**Figure S7. (a)** Diagrammatic representation of all significantly impacted [increased: blue, decreased: red] pantothenate and CoA metabolites in MRSA ATCC 43300 following treatment with C12-o-(BG-D)-10. **(b)** Significantly impacted pantothenate and CoA metabolites in MRSA ATCC 43300 following treatment with C12-o-(BG-D)-10 at 1 h (red), 3 h (green), and 6 h (blue). Putative metabolite names are assigned based on accurate mass (≥1.0-log2-FC; $p$<0.05).

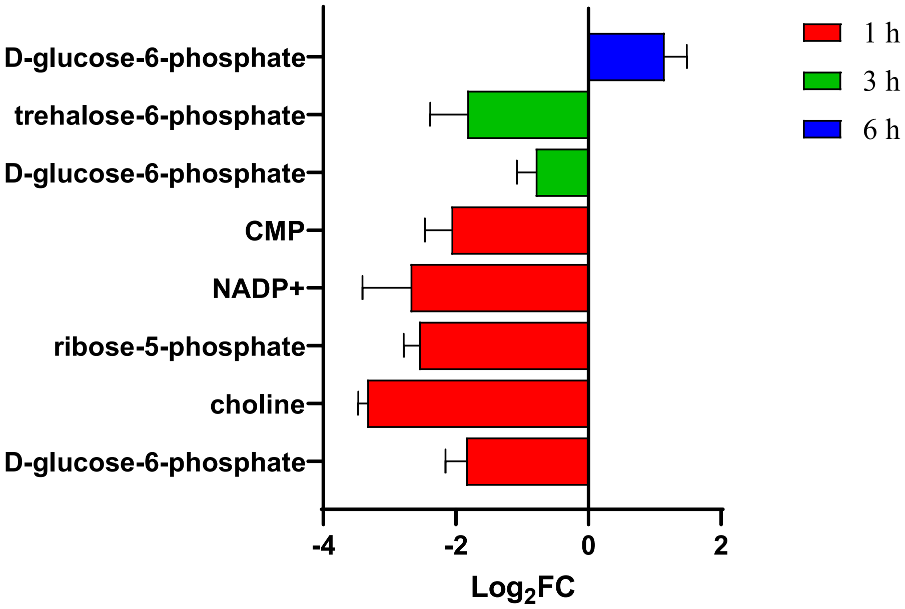


**Figure S8.** Significantly impacted homeostasis and stress pathways metabolites in MRSA ATCC 43300 following treatment with $C_{12}$-o-(BG-D)-10 at 1 h (red), 3 h (green), and 6 h (blue). Putative metabolite names are assigned based on accurate mass (≥1.0-$\log_2$-FC; $p$<0.05).

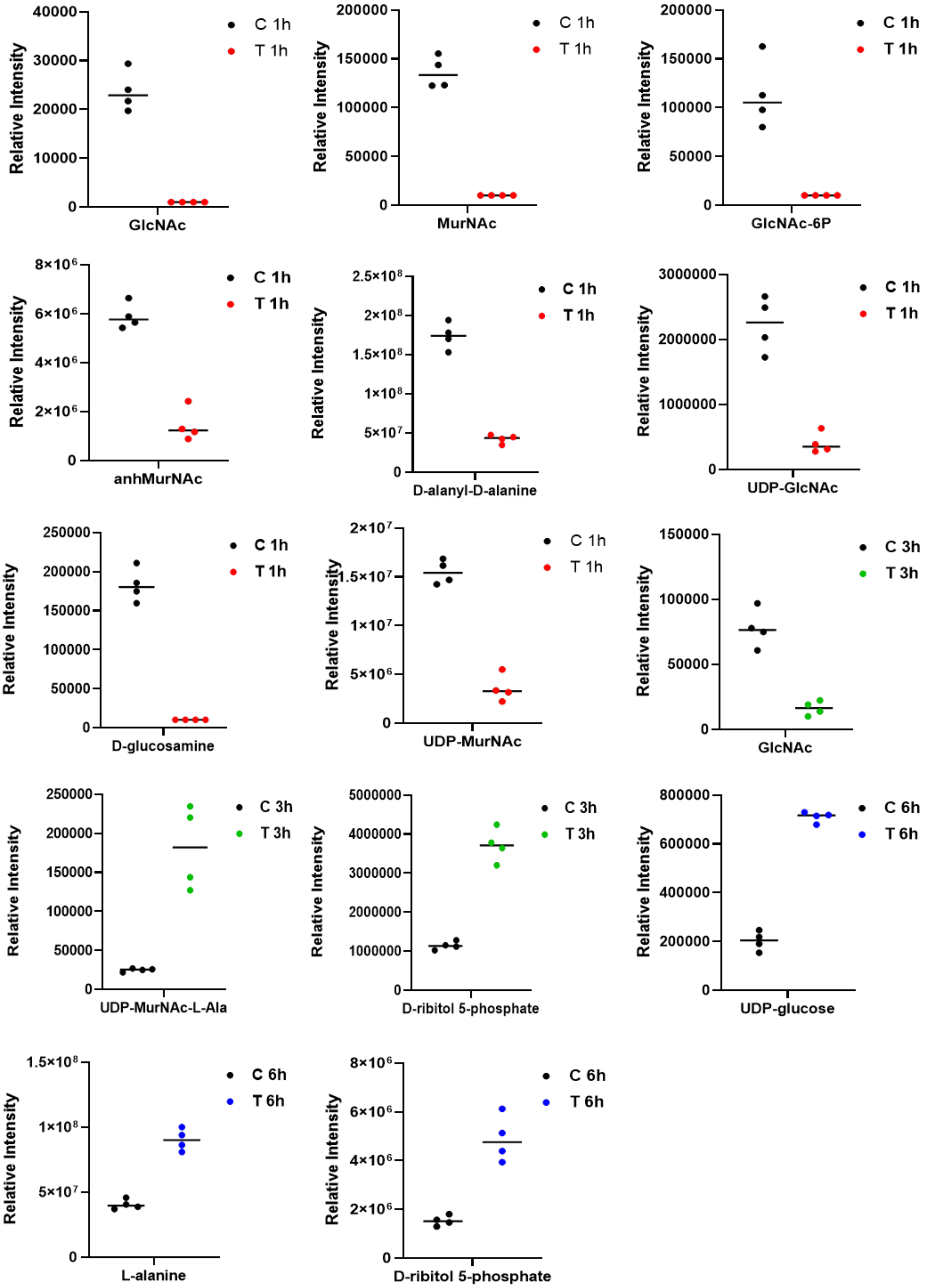


**Figure S9.** Individual values plot for significantly perturbed amino-sugar and sugar-nucleotide metabolites of MRSA ATCC 43300 following $C_{12}$-o-(BG-D)-10 treatment at 1h (red), 3h (green), and 6h (blue) (≥ 1.0-$\log_2$-FC; $P<0.05$). C, control (untreated); T, $C_{12}$-o-(BG-D)-10.

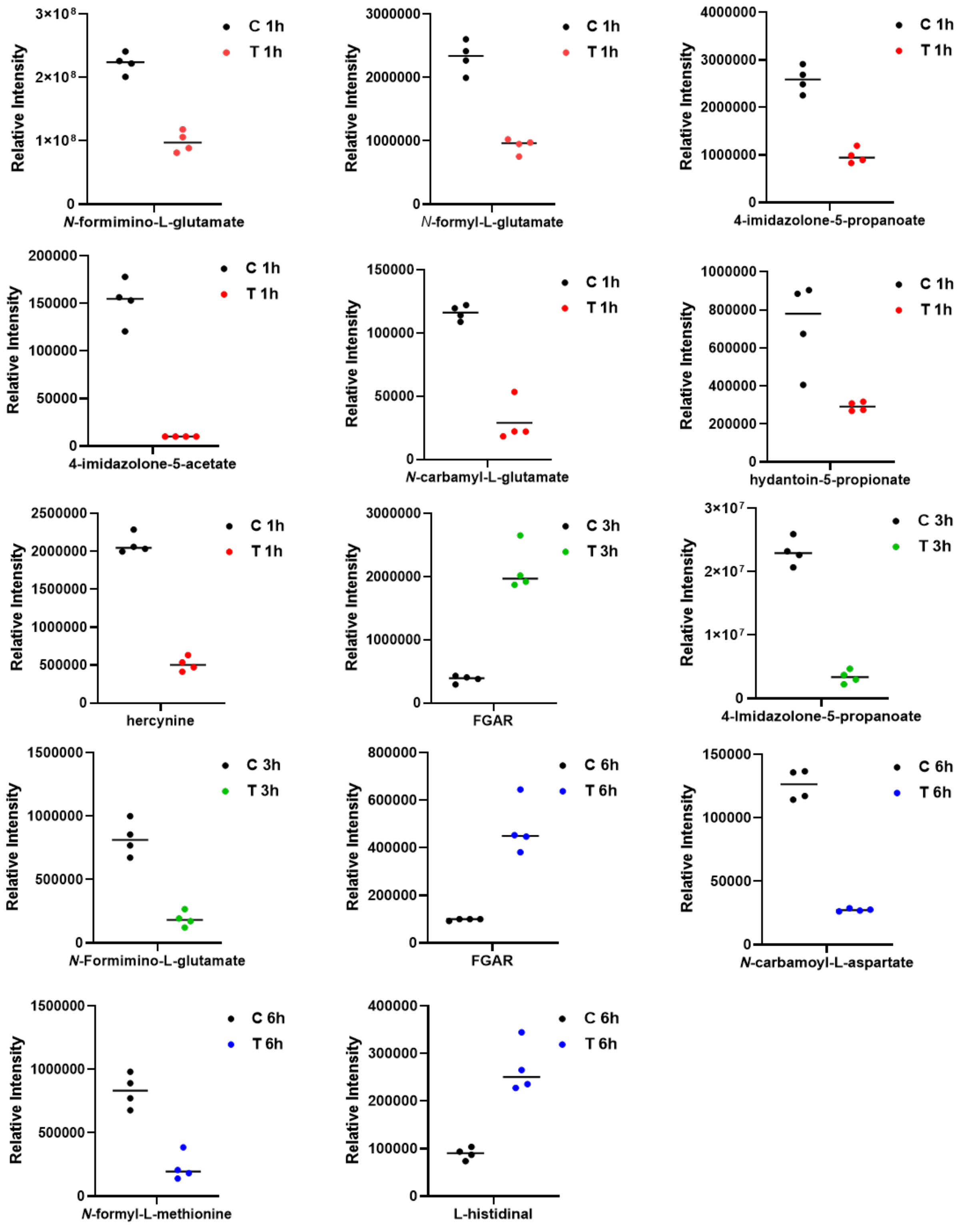


**Figure S10.** Individual values plot for significantly perturbed histidine metabolites of MRSA ATCC 43300 following $C_{12}$-o-(BG-D)-10 treatment at 1h (red), 3h (green), and 6h (blue) (≥ 1.0-$\log_2$-FC; $P$<0.05). C, control (untreated); T, $C_{12}$-o-(BG-D)-10.

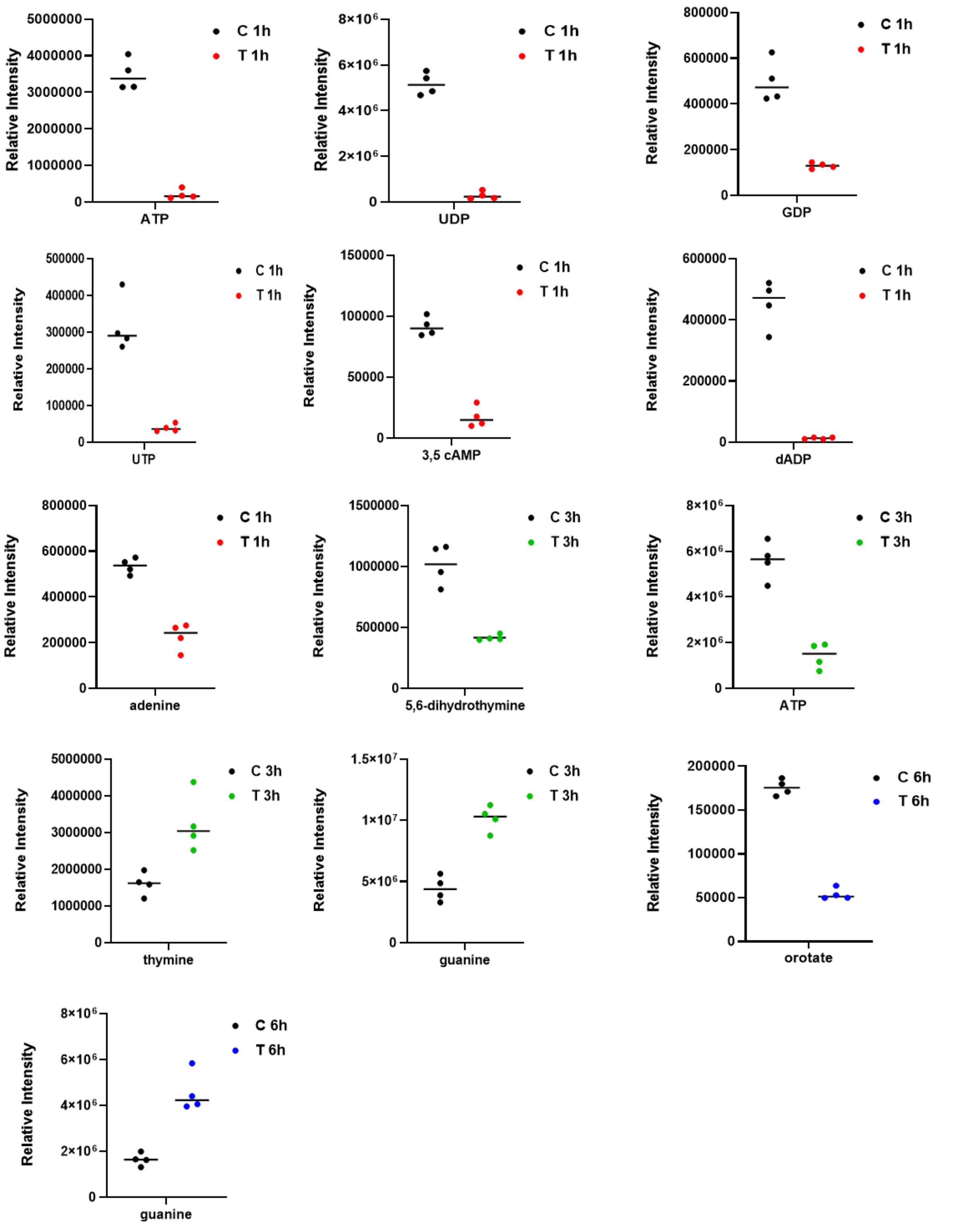


**Figure S11.** Individual values plot for significantly perturbed pyrimidine and purine metabolites of MRSA ATCC 43300 following $C_{12}$-o-(BG-D)-10 treatment at 1h (red), 3h (green), and 6h (blue) (≥ 1.0-$log_2$-FC; $P<0.05$). C, control (untreated); T, $C_{12}$-o-(BG-D)-10.

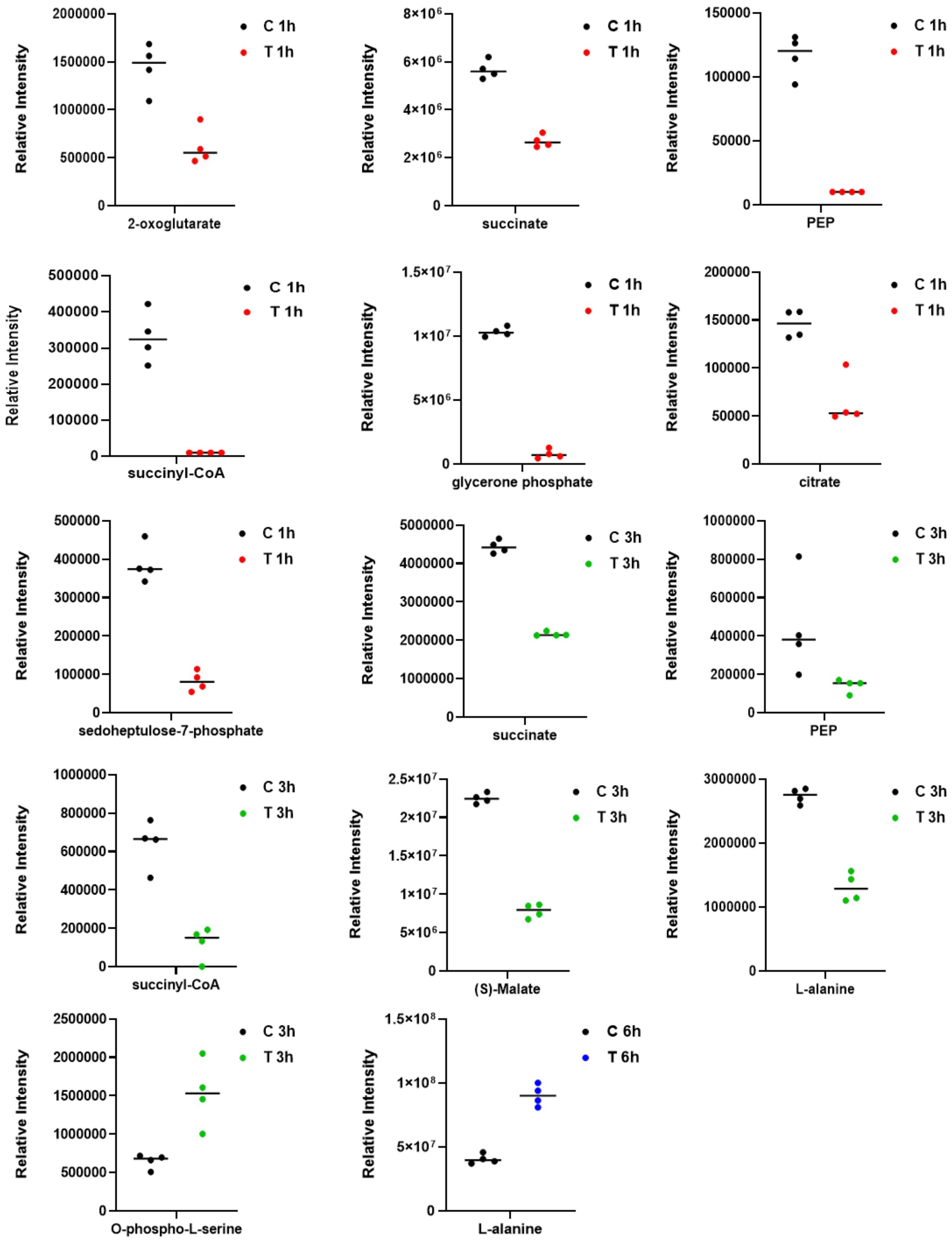


**Figure S12.** Individual values plot for significantly perturbed central carbon metabolism metabolites of MRSA ATCC 43300 following $C_{12}$-o-(BG-D)-10 treatment at 1h (red), 3h (green), and 6h (blue) ($\geq$ 1.0-$log_2$-FC; $P$<0.05). C, control (untreated); T, $C_{12}$-o-(BG-D)-10.

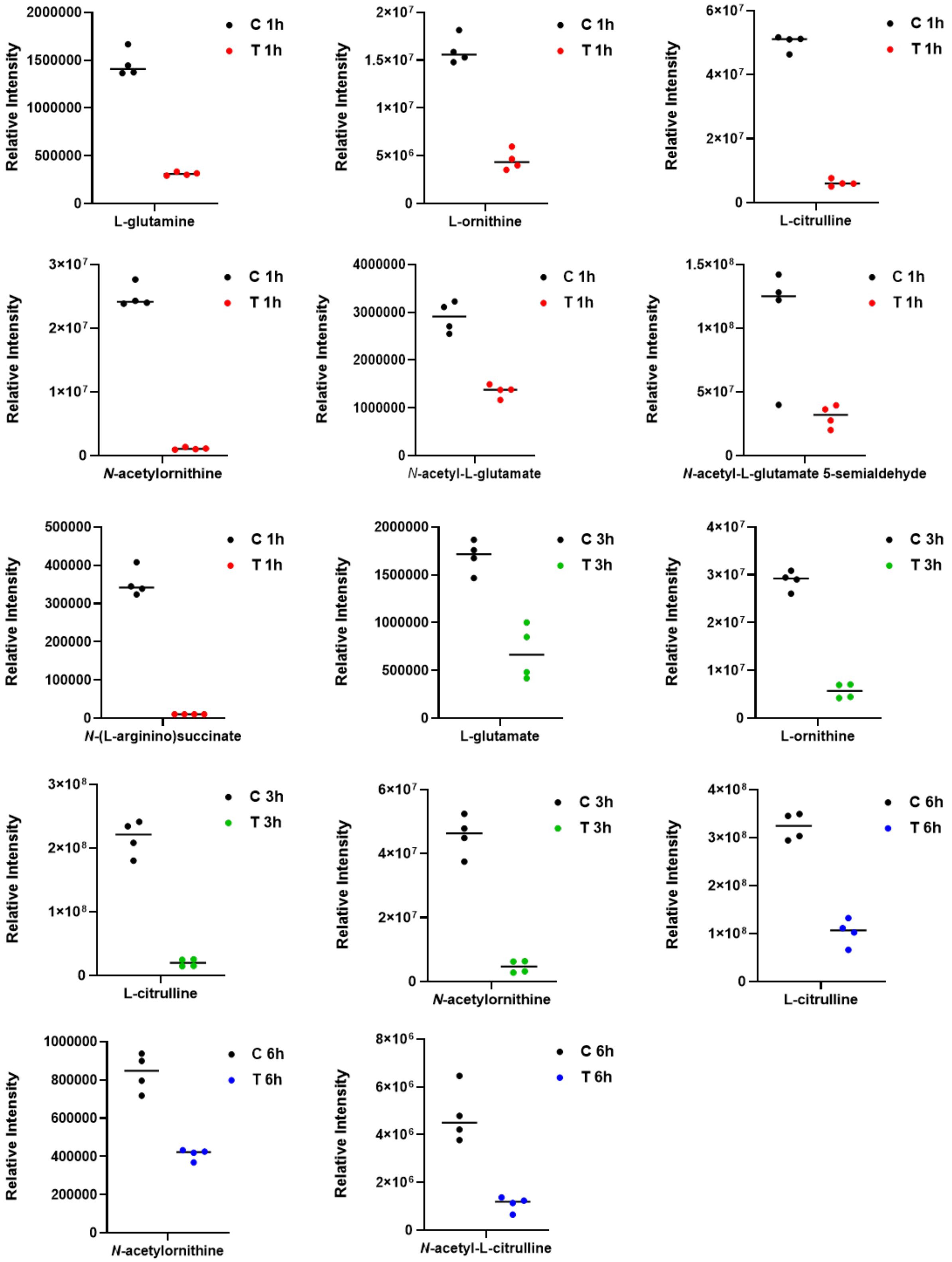


**Figure S13.** Individual values plot for significantly perturbed arginine and interrelated TCA cycle metabolites of MRSA ATCC 43300 following $C_{12}$-o-(BG-D)-10 treatment at 1h (red), 3h (green), and 6h (blue) (≥ 1.0-$log_2$-fold; $P$<0.05). C, control (untreated); T, $C_{12}$-o-(BG-D)-10.

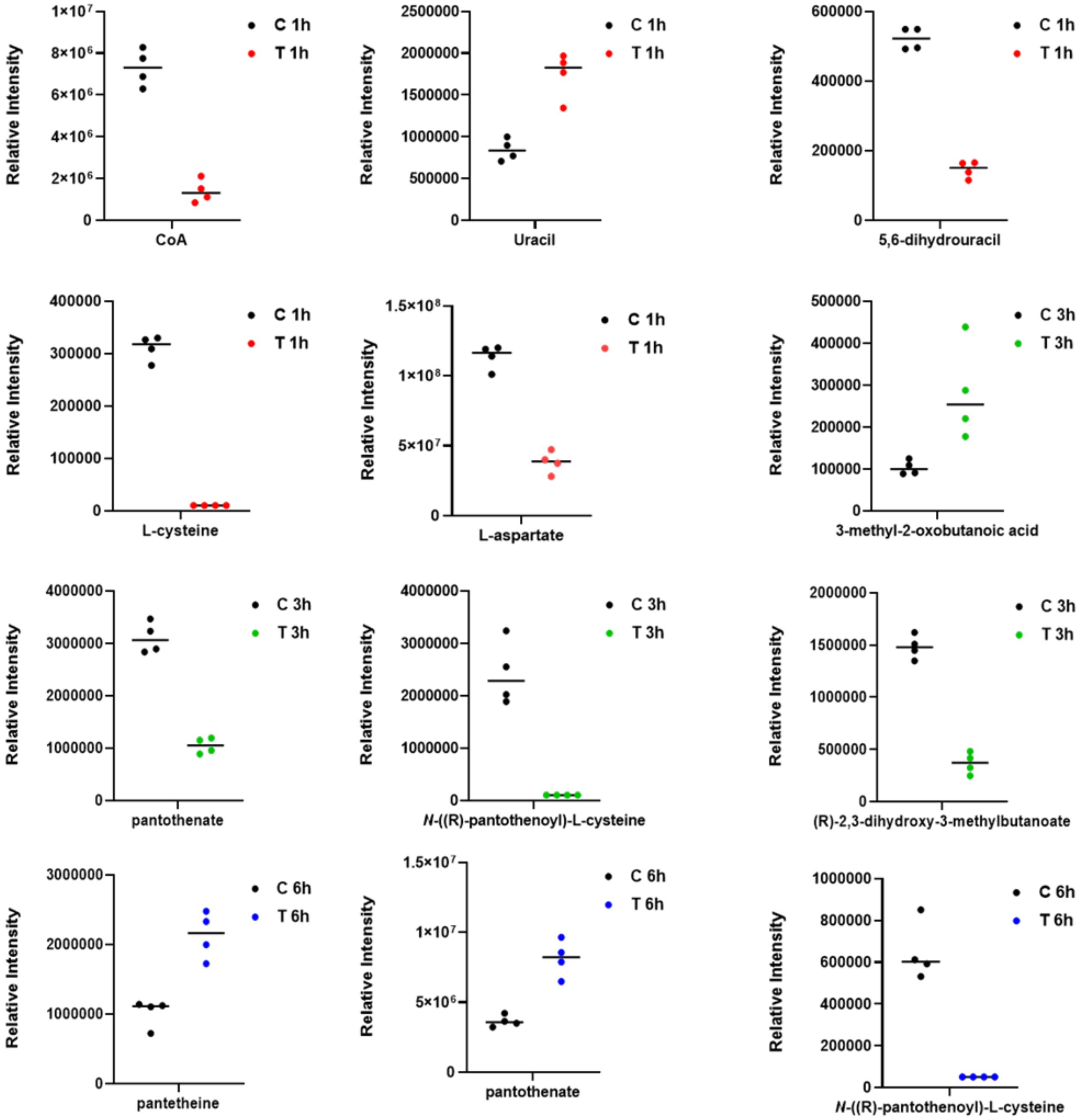


**Figure S14.** Individual values plot for significantly perturbed pantothenate and CoA metabolites of MRSA ATCC 43300 following $C_{12}$-o-(BG-D)-10 treatment at 1h (red), 3h (green), and 6h (blue) ($\geq$ 1.0-$\log_2$-fold; *P*<0.05). C, control (untreated); T, $C_{12}$-o-(BG-D)-10.
